# Nanoscale Optical Imaging, Reconstruction, and Spatial Analysis of Whole Mouse Glomeruli

**DOI:** 10.1101/2024.10.31.620364

**Authors:** Adilijiang Ali, Zixuan Liu, Kenan Ye, Yun Guan, Siying Chen, Tingxuan Liu, Ziyu Guo, Madeline K. Wong, Pedro Vasquez, Chetan Poudel, Benjamin C. Mustonen, Diana G. Eng, Jeffrey W. Pippin, Stuart J. Shankland, Sheng Wang, Joshua C. Vaughan

## Abstract

Renal glomeruli have traditionally been studied by micrometer-scale optical microscopy to interrogate overall physiology or molecular distributions and by nanoscale electron microscopy to interrogate the ultrastructure of thin sections. While these approaches are powerful, they have been limited in their ability to obtain detailed views of the glomeruli as holistic 3D functional units. To fill this knowledge gap, we have developed a novel pipeline for imaging, reconstructing, and analyzing whole mouse glomeruli at 100 nm resolution using super-resolution fluorescence microscopy. This pipeline integrates both manual and machine learning approaches to annotate and analyze glomerular structures. Using this method, we created 18 detailed glomerulus models, from a range of healthy, aged, and model diseased mice, that outline all major structures and cell types. These models have been made publicly accessible in an online repository, providing a valuable resource for further studies. Our results also uncovered a diverse set of novel phenotypes including nuclear enlargement in all glomerular cell types in aging and disease, as well as an aging-related pattern of regional thickening of the Bowman’s capsule basement membrane near the tubular-glomerular junction.

## Introduction

The primary function of renal glomeruli is to filter the blood. This requires an anatomically complex makeup comprising capillary loops, several different resident cell types, and basement membranes. Glomerular function diminishes with age and disease, leading to impaired kidney function and serious deterioration of health. Much of our understanding of glomeruli, and the kidney overall, derives from microscopy studies in two distinct regimes. Optical microscopy provides micron-scale snapshots of overall physiology and distributions of key molecules (e.g., immune complexes) ^1–9^ and electron microscopy (EM) provides detailed nanometer-scale views of glomerular ultrastructure ^10–14^. These traditional methods, while powerful, have faced several shortcomings. First, they are by necessity technically performed using different pieces of tissue and fixatives, preventing each method from being directly correlated. This has the potential to produce contradictory results. Second, high-resolution measurements of glomeruli are generally taken using EM to measure two-dimensional thin sections at random angles or three-dimensional analysis of subregions (e.g., serial block face EM). This can create challenges with interpretation and sometimes overlook critical details. These limitations have constrained research and clinical practice to focus on specific subsets of structures or small regions of the glomerulus, rather than allowing for a comprehensive study of the entire glomerulus as a functioning unit.

To close these technical and knowledge gaps, we have used super-resolution optical microscopy and a computational analysis pipeline to create a shared resource consisting of high-resolution (∼100 nm) annotated data sets of whole mouse glomeruli for healthy, aged, and disease model tissue. The datasets provide detailed 3D models with annotations of all major structural components of glomeruli and renal corpuscle (i.e., Bowman’s capsule basement membrane (BCBM), glomerular basement membrane (GBM), capillary lumen, and Bowman’s space) as well as cell number and locations for all primary glomerular cell types (podocytes, mesangial cells (MCs), glomerular endothelial cells (GEnCs), and parietal epithelial cells (PECs)). The shared data repository is a free, publicly accessible digital resource that can be mined by nephrology, pathology, computational, and other communities for diverse purposes.

## Methods

### Mouse tissue

C57BL/6 mice were used for healthy, aged, and D14 FSGS mouse models. Aged mice from 27-29 m.o. (∼75+ years old human) were acquired from the National Institute of Aging. For the podocyte depletion FSGS model, mice were given 2 consecutive doses of anti-podocyte antibody at 10mg/20gbw and euthanized after 14 days. Anesthetized adult mice were cardiac perfused with PBS/Azide (phosphate buffered saline containing 3 mM sodium azide) and 4% paraformaldehyde (PFA), each for 4.5 minutes. Both kidneys were harvested, and the renal capsules were removed. Each kidney was then fixed in 10 ml of the 4% PBS/PFA solution for 1 hour. All animal protocols (4442-01, 2968-04) were approved by the University of Washington Institutional Animal Care and Use Committee (IACUC). After fixation, the kidney was embedded in 4% (w/v) agarose in 1× PBS/Azide and then sliced into 100 µm sections using a vibratome.

### Expansion microscopy

Mouse tissue sections were gelled, denatured, and expanded to achieve ∼4× isotropic expansion in all directions as described in the **Supplementary Methods**.

### FLARE staining protocol

Tissues were stained for carbohydrates, amines, and DNA using the FLARE protocol as described in the **Supplementary Methods**.

### Fluorescence Image Acquisition

All images were recorded using a scanning confocal microscope (Nikon A1R HD25 laser scanning confocal microscope, LSB B202A), with a 40× water immersion objective lens (Apochromat Lambda S LWD 40× water, NA 1.15, WD 0.59–0.61 mm).

### Glomerular structures segmentation

All the imaged stacks were segmented using a combination of manual and semiautomated methods as described in the **Supplementary Methods**.

### Glomerular structures reconstruction and analysis

The segmented results were reconstructed using Imaris (RRID: SCR_007370). For specific instructions on the surface creation tool, see Imaris Reference Manual (https://imaris.oxinst.com/).

### Basement membrane thickness analysis

Basement membrane thickness was measured using a custom MATLAB script (https://doi.org/10.5281/zenodo.13146131). In general, segmented masks were first scaled to ensure isotropic voxels. These scaled masks were used to generate surface normal vectors for each vertex. A line centered at each vertex was used to measure the intensity profile of the raw image stack along the direction of the corresponding surface normal vector. A Gaussian function was fitted to each intensity profile, and the thickness was estimated from the FWHM of the fitted Gaussian curve.

## Data availability

All imaging, model datasets, and detailed handling instructions are available on the SharePoint repository (https://uwnetid.sharepoint.com/:f:/s/mouse_glomerulus_repository/EtSJNi1PmudNrhZMUdFtBD8BHWRcrg3×0NwSs7AZXTOJKA?e=NyKA9O).

## Results

### High-resolution, 3D fluorescence imaging of whole glomeruli

The pipeline for 100 nm resolution imaging, segmentation, and analysis of whole mouse glomeruli is shown in **Figure 1**. The pipeline uses expansion microscopy, a technique that makes small samples physically larger by embedding and expanding them in a swellable hydrogel to enable high-resolution imaging^15–17^ and also employs a recently developed tri-color fluorescent stain called FLARE to stain general kidney structure. FLARE is a fluorescent analog of a combined hematoxylin and eosin (H&E) and periodic acid Schiff (PAS) stain^18,19^ that labels nuclei (DNA, blue), proteins (primary amines, red), and carbohydrates (green). Confocal microscopy was used to acquire three-channel data stacks of whole glomeruli within expanded mouse kidney tissue sections, where data sets consisted of about 2048×2048×400 voxels per glomerulus. Only cortical glomeruli were studied since complete juxtamedullary glomeruli were typically larger than our 100 μm tissue sections and larger than our microscope’s field of view (**Supplementary Figure S1**).

**Figure 1.**
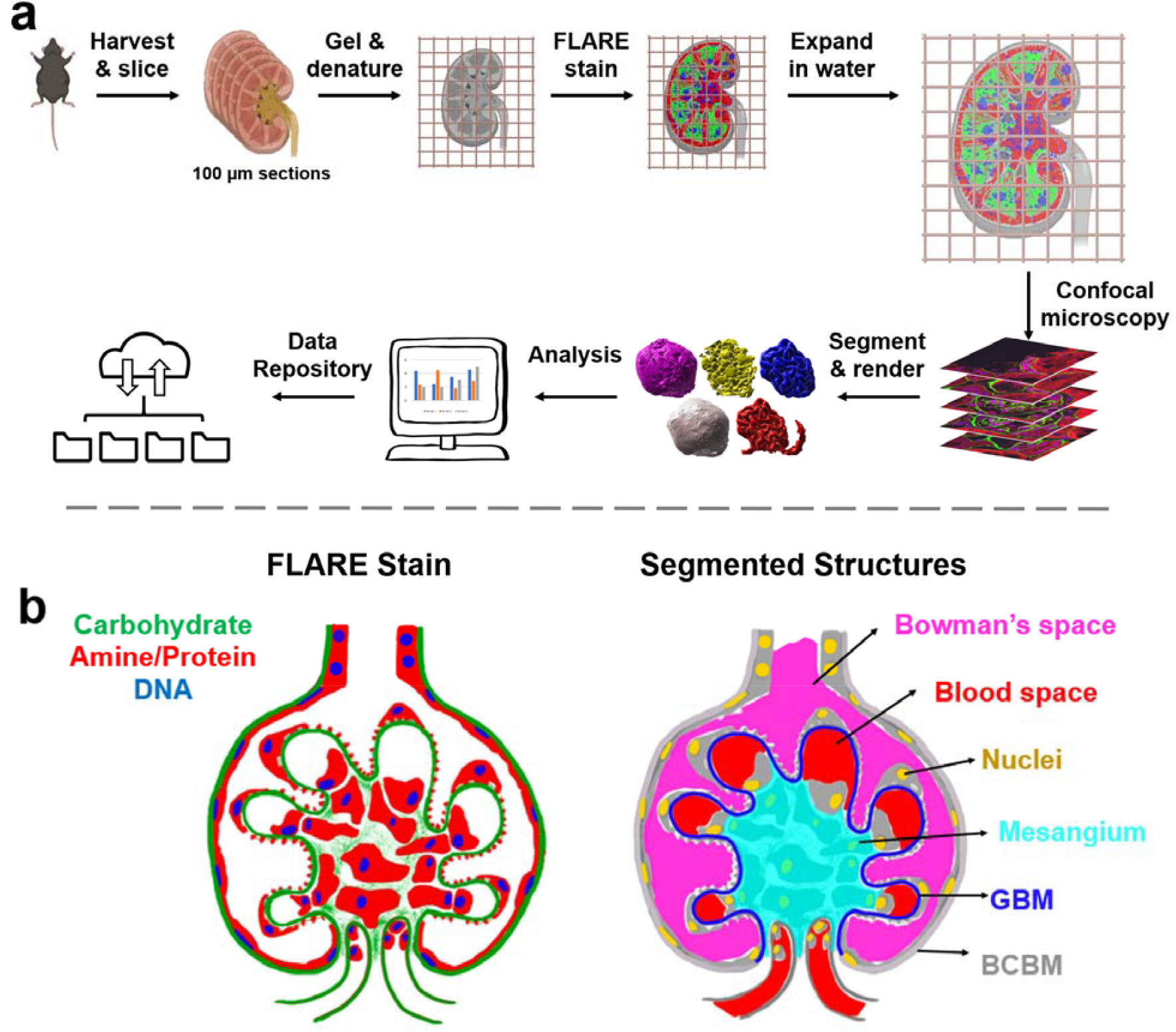
Schematic illustration of a) the project workflow and b) segmented glomerular structures. In b), the left half shows a FLARE-stained glomerulus while the right half demonstrates how glomerular structures are represented based on the FLARE-stained sample. Bowman’s space and the blood space are defined as empty spaces where there is no FLARE signal, with Bowman’s space outside the GBM but within the Bowman’s capsule and blood space inside the GBM. The BCBM and GBM are differentiated at the point where the carbohydrate signal at the periphery invaginates towards the inner glomerular space. The color schemes in b) are consistent with all raw data and segmented & reconstructed volumes in this report.

The resulting three-channel image stacks revealed key structural components of the glomerulus and its surrounding structures (**Figure 2**, left half of each panel; **Supplementary Movie S1-3**). A serial section of a FLARE-stained and expanded mouse glomerulus revealed diverse glomerular structures, including the carbohydrate-rich BCBM, GBM, and proximal tubular basement membrane (**Supplementary Figure S2a-f**). Furthermore, finer structures, such as interdigitated podocyte foot processes (**Supplementary Figure S2g, h**), and nuclear chromatin organization (**Supplementary Figure S2i**) were visible.

**Figure 2.**
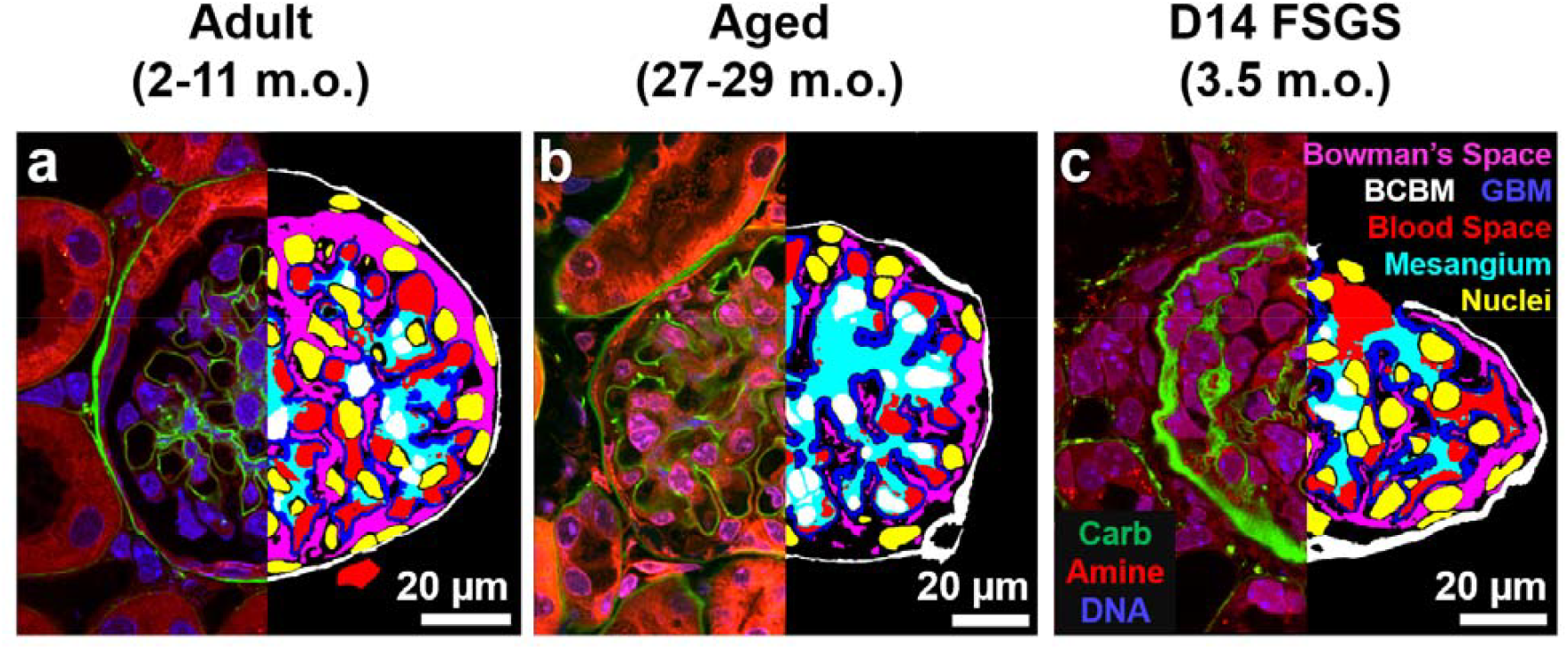
Representative single planes from volumetric raw data (left) and segmented structures (right) for cortical a) adult, b) aged, and c) D14 FSGS glomeruli. Left halves: raw three-channel FLARE image. Green = carbohydrate, red = amine/protein, blue = nuclei/DNA. Right halves: segmentation results. Red = blood space, blue = GBM & mesangium, yellow = nuclei (the white color in the middle is due to the overlap with cyan), white = BCBM, and magenta = Bowman’s space. Scale bars: 20 µm in pre-expansion units.

### Segmentation and reconstruction of mouse glomeruli

The bulk of the labor of the pipeline was devoted to computational analysis of the confocal microscopy data to segment out the six key structural components from the volumetric data, including the BCBM, GBM, blood space, mesangium, Bowman’s space, and cell nuclei. This challenge was addressed in two phases. In the first phase, we used fully manual image segmentation (i.e., tracing images on a tablet) to segment whole glomeruli. Given the size and complexity of the multichannel, volumetric microscopy data sets, manual segmentation of each glomerulus required ∼4 weeks of analysis. In the second phase, we used the first 5 analyzed glomeruli and the raw data sets to train a machine learning algorithm (**Supplementary Figure S3**) to make good initial predictions for the six main structural components. Subsequent data sets were initially segmented using the trained model and then manually reviewed and/or corrected. This hybrid approach required considerably fewer human hours, at ∼3 days of human effort per glomerulus.

Each of the six structural components of the glomerulus was defined as illustrated in **Figure 1b** and segmented as described in the **Supplementary Methods**. All the segmented results were further processed for 3D reconstruction using Imaris. The representations of a single cortical glomerulus from healthy adult, aged, and diseased samples are shown in **Figure 3 a-b**. All structures are oriented with capillary poles at the bottom and urinary poles at the top for easy comparison. The comprehensive collection of the entire set of glomeruli is presented in **Supplementary Figure S4**.

**Figure 3.**
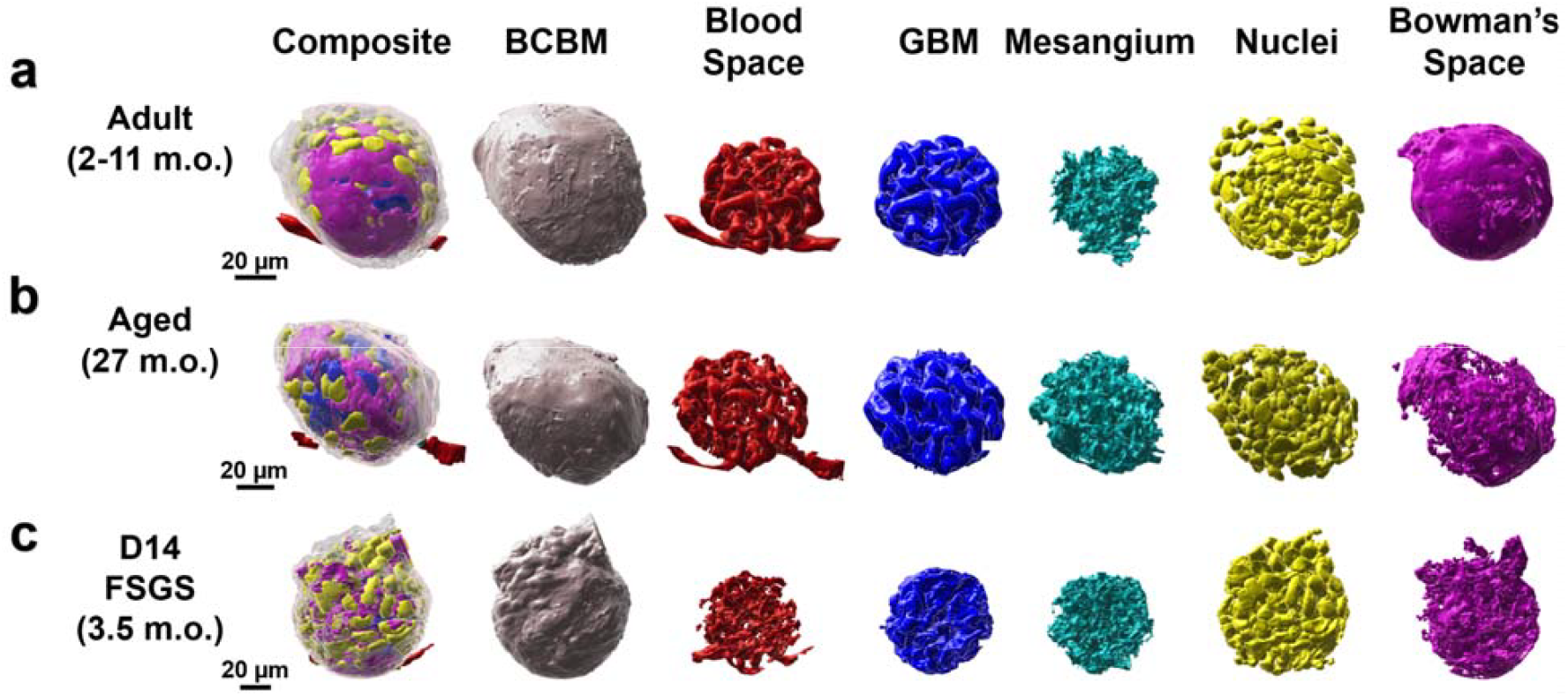
Reconstructed cortical glomeruli for a) adult, b) aged, and c) day 14 FSGS mice. All the glomeruli are oriented with their capillary poles at the bottom and their urinary poles at the top. All scale bars are in pre-expansion units.

### Whole-glomerulus morphometric quantification

The reconstruction of whole glomeruli enabled the unique, holistic analysis of glomerular properties rather than limiting sampling to small regions of individual glomeruli or thin sections sampled at random angles which can suffer projection artifacts and comparable quantification across glomeruli from adult, aged, and experimental focal segmental glomerulosclerosis (FSGS) mice.

We observed notable differences in the overall volume of the renal corpuscles (**Figure 4a**) and the volume of key glomerular structures (**Figure 4b**). The 2-11 m.o. healthy group had an overall volume of the renal corpuscle of 4 ± 1.4×10^5^ μm^3^ (mean ± standard deviation). In the aged mouse kidney at 27-29 months of age, the renal corpuscle volume was ∼2-fold larger, wherea the FSGS group was only slightly larger. More strikingly, the aged and FSGS model system showed dramatic increases in the mesangial size, both in absolute volume and as a fraction of the renal corpuscle volume (**Figure 4b-c**). The enlargement of the mesangium was accompanied by reductions in both blood space and Bowman’s space, consistent with a prior report for aged and FSGS where mesangial expansion often correlates with capillary collapse and diminished urinary space^20^.

**Figure 4.**
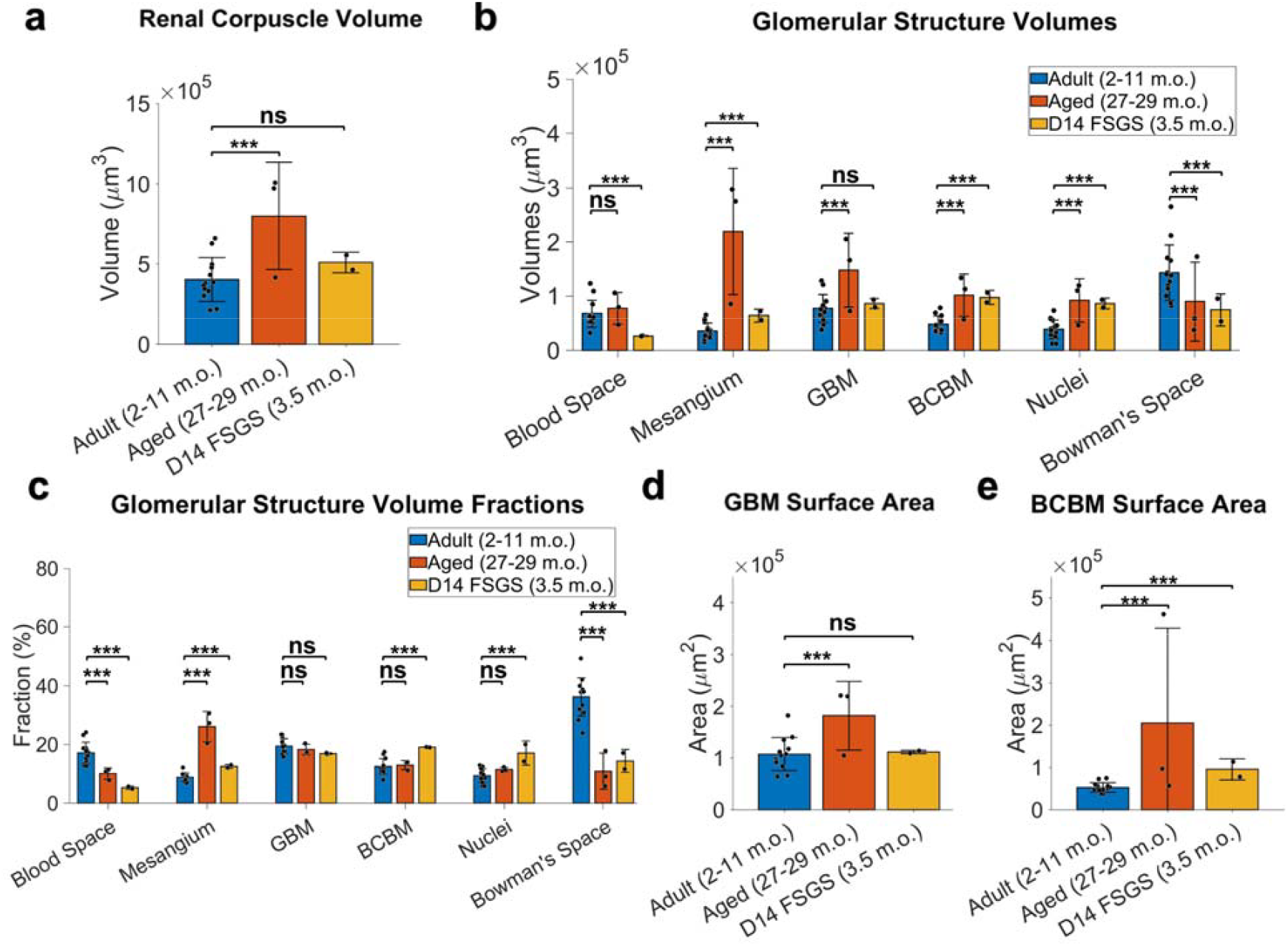
Whole glomerulus level quantitation of glomerular structures. a) Renal corpuscle volumes. b) volumes of individual structures. c) Relative proportions of glomerular structure volume to the renal corpuscle volume. d) GBM surface area. e) BCBM surface area. All the measurements are in pre-expansion units. Error bars indicate mean ± standard deviation. *** p < 0.05, ns: not significant.

A distinct feature of FSGS is PEC activation and their movement towards the sites of podocyte loss on the GBM, depositing matrix along the way, expanding the BCBM into the inner glomerular space^21,22^. Consistent with this picture, we observed an increased BCBM volume fraction in FSGS from 12.5 ± 2.6% to 19.1 ± 0.1% (**Figure 4c**).

From our models, we calculated the total surface area of the GBM to be 1.1 ± 0.3×10^5^ μm^2^ for the healthy adult and FSGS glomeruli (**Supplementary Table S1-2**) but was nearly double that for the aged kidney (**Figure 4d**). The total surface area of the BCBM was significantly increased in both aged and FSGS kidneys (**Figure 4e**). For reference, an area of 1.1×10^5^ μm^2^ corresponds to the surface area of a sphere with a radius of ∼94 μm.

### Glomerular cell type classification and cell morphometry

The detailed high-resolution views of the whole glomeruli enabled the classification of all glomerular cell types based on their nuclear shape and position relative to other glomerular structures **(Figure 5a, Supplementary Movie S4**). An example classification frame of all cell types is presented in **Supplementary Figure S6**: PEC nuclei are relatively flat and positioned along the BCBM, podocyte nuclei are along the GBM but face the urinary space, mesangial nuclei are contained within the mesangium without facing the blood space, and GEnC nuclei face the blood space. We performed independent validation experiments using five established cell-type specific antibodies ^23–28^ (**Supplementary Figure S7**) and determined that the cell classification procedure was highly accurate for all cell types with precision and recall values of 91-100% (**Supplementary Table S3**). Although some immune cells are also expected, the number was evidently small. By this means, detailed maps of all cell nucleus positions within the glomeruli were obtained.

**Figure 5.**
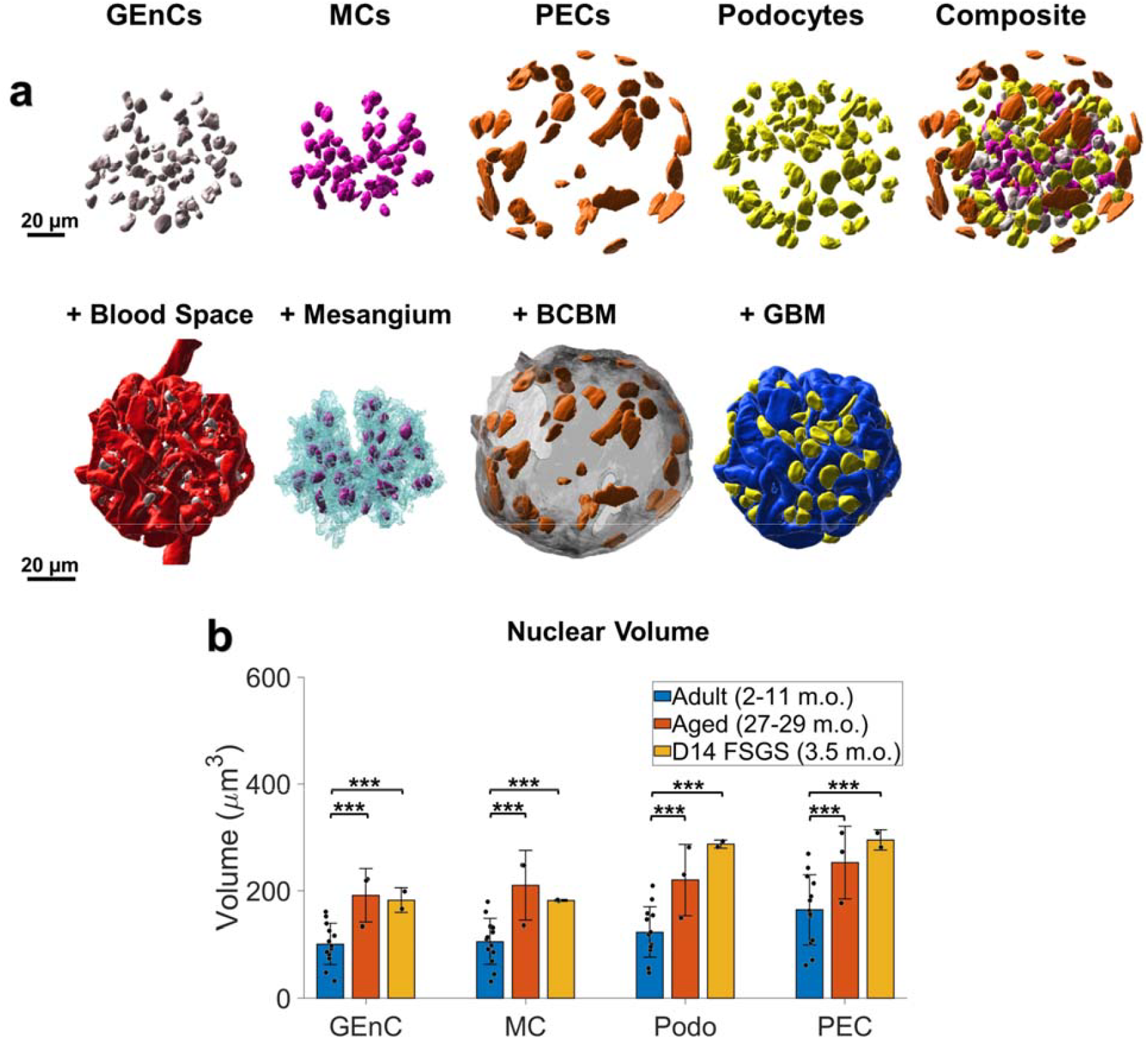
Whole glomerulus cell type classification and analysis. a) An example of one healthy adult mouse glomerulus with reconstructed nuclei of individual cell types (top row) the relative positioning of nuclei and their glomerular compartments (bottom row). b) Comparison among glomerular cell nuclei volume in healthy adults, aged, and FSGS. Scale bars = 20 μm. All measurements are in pre-expansion units. *** p < 0.05, ns: not significant.

Our validated cell type analysis revealed an average of 262 ± 52 glomerular cells (GEnCs + MCs + podocytes) in the glomerular tuft of healthy adult mice (**Table 1**). This is slightly higher than the previously reported value of 210-235 cells per glomerulus^29^. This discrepancy may result from a combination of factors, including the higher spatial resolution of our approach compared to traditional optical microscopy methods or differences in ages, strains, or sampling. Notably, the total glomerular cells were increased 33% in aged mouse samples compared to healthy adult mice, largely as a result of more abundant GEnCs and MCs (**Table 1**).

**Table 1.** The number & relative fraction of glomerular cell types indicated as mean (± standard deviation).

|  | Adult (3-15 m.o.) |  | Aged (27-29 m.o.) |  | D14 FSGS (3.5 m.o.) |  |
| --- | --- | --- | --- | --- | --- | --- |
|  | Count | Fraction | Count | Fraction | Count | Fraction |
| **Glomerular Endothelial Cell** | 92<br>($\pm 26$) | 28.5%<br>($\pm 3.8\%$) | 152<br>($\pm 53$) | 35.2%<br>($\pm 6.1\%$) | 106<br>($\pm 7$) | 29.2%<br>($\pm 1.6\%$) |
| **Mesangial Cell** | 73<br>($\pm 14$) | 22.8%<br>($\pm 1.9\%$) | 133<br>($\pm 44$) | 31.0%<br>($\pm 5.4\%$) | 85<br>($\pm 19$) | 23.1%<br>($\pm 2.5\%$) |
| **Podocyte** | 97<br>($\pm 22$) | 30.7%<br>($\pm 5.5\%$) | 91<br>($\pm 15$) | 22.5%<br>($\pm 7.1\%$) | 54<br>($\pm 25$) | 14.4%<br>($\pm 5.1\%$) |
| **Parietal Epithelial Cell** | 58<br>($\pm 16$) | 18.1%<br>($\pm 4.3$) | 46<br>($\pm 2$) | 11.3%<br>($\pm 2.4\%$) | 120<br>($\pm 7$) | 33.3%<br>($\pm 6.0\%$) |
| **Total** | 319<br>($\pm 60$) | 100% | 423<br>($\pm 87$) | 100% | 364<br>($\pm 44$) | 100% |

### Cell type analysis

In healthy kidney samples, an average of 58 ± 16 PECs were identified in healthy adult samples, comprising 18.1 ± 4.3% of total glomerular cells (**Table 1**). There were no significant changes in PEC counts in aged kidneys, although the percentage of PECs decreased substantially due to an increase in GEnCs and MCs. The nuclei of PECs consistently exhibited a flatter shape compared to the other cell types (**Figure 5a, Supplementary Movie S4**).

In a state of podocyte depletion, PECs serve as potential progenitor cells to generate new podocytes^30–32^. Activated PECs migrate toward the glomerular tuft and deposit extracellular matrix proteins underlying the development of glomerulosclerosis. Both the number and the relative proportions of PECs nearly doubled in mice with FSGS (**Table 1**).

An average of 97 ± 22 podocytes per glomerulus were identified in healthy adult mice. Podocyte number decreased in FSGS (**Table 1**) as expected based on previous reports. No significant changes in podocyte number were observed in aged kidneys, likely due to the limited sample size. However, the relative proportion of podocytes decreased notably in both FSGS and the aged kidney. These results are consistent with previous reports in aging and in FSGS^20,33,34^.

Healthy adult samples had 92 ± 26 GECs and 73 ± 14 MCs, composing 28.5 ± 3.8% and 22.8 ± 1.9% of cells, respectively (**Table 1**). There was a notable increase in the number and relative proportion of GECs and MCs in aged kidneys. Although there was an overall increase in GEC and MC counts in FSGS samples, the relative proportions remained unchanged.

It has been reported that the size of podocyte nuclei increases by hypertrophy as a compensation mechanism for the depleted podocyte counts^20^. Indeed, we found that the average podocyte nuclear volume increased significantly in both the aged and FSGS kidneys (**Figure 5b**). The results showed that the nuclei of all four cell types (GEC, MC, PEC, and podocyte) were also significantly larger in aged samples compared to young healthy kidneys (**Figure 5b**).

### Whole-glomerulus quantitation of basement membrane thickness

We used our high-resolution reconstructions to perform global analysis of the thickness distribution of both the BCBM and the GBM for the first time in whole glomeruli. Membrane thickness was quantified by first calculating the carbohydrate channel’s intensity profile perpendicular to each basement membrane at tens to hundreds of thousands of positions across the BCBM or GBM of each glomerulus and then fitting each intensity profile with a Gaussian function to determine the full width at half maximum as a measure of thickness (FWHM, **Figure 6a**).

**Figure 6.**
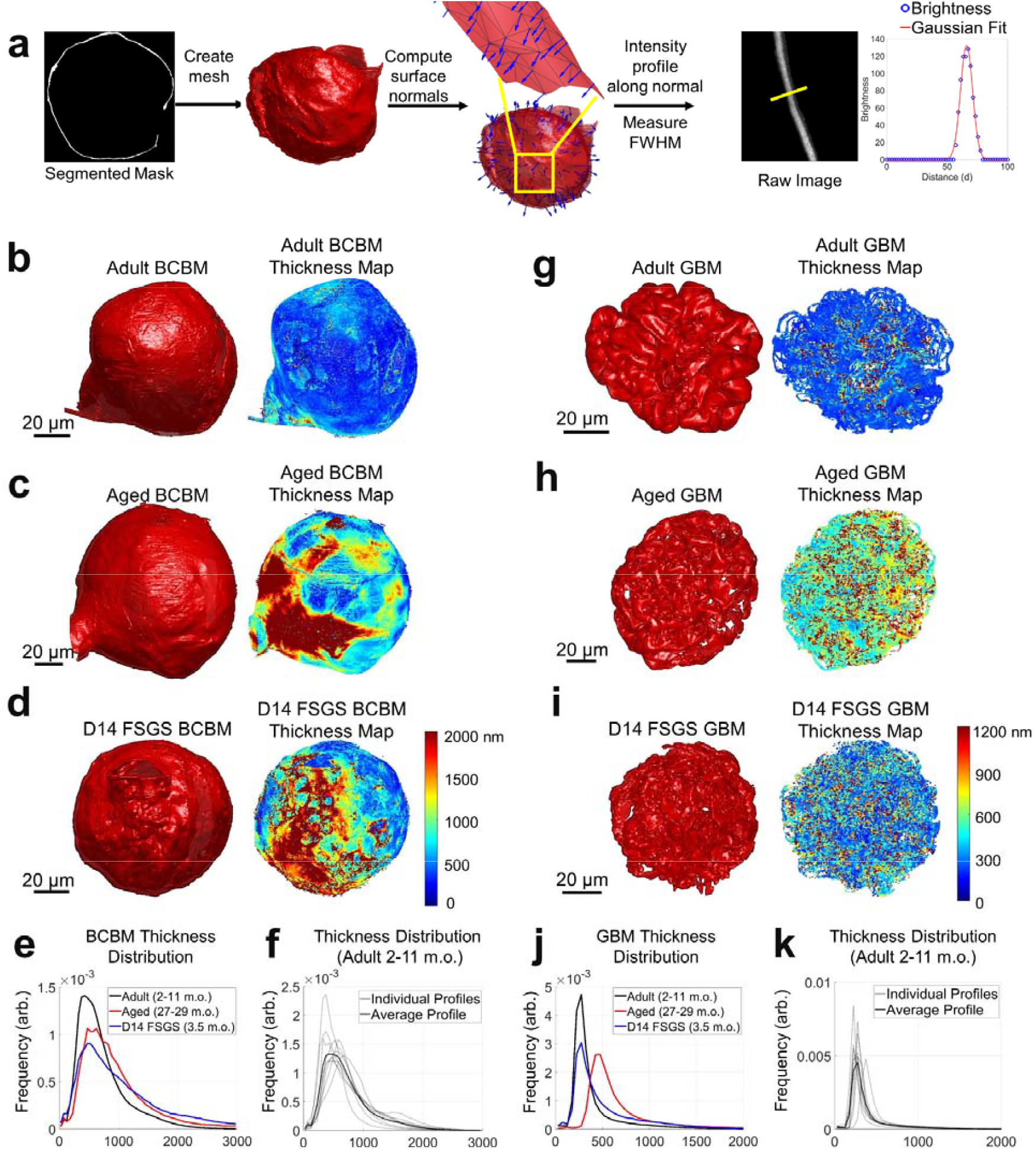
Thickness distribution analysis of the Bowman’s capsule basement membrane (BCBM) and glomerular basement membrane (GBM). **a)** Schematic illustration of the thickness analysis process: Meshes were generated from segmented masks of the BCBM/GBM. Surface normal vectors were calculated for each mesh face, and intensity profiles were extracted along these normal vectors from the raw image data. A Gaussian function was fitted to each profile, and thickness was estimated using the full width at half maximum (FWHM) of the Gaussian curve. **b-d)** 3D meshes and corresponding thickness maps of BCBM of: **b)** adult (2–11 m.o.), **c)** aged (27– 29 m.o.), and **d)** D14 FSGS (2–11 m.o.) glomerulus. **e)** Line profiles of the thickness distribution of BCBM for adult, aged, and D14 FSGS glomeruli. **f)** Average and individual thickness distributions of adult BCBM. **g-i)** 3D meshes and corresponding thickness maps of GBM: **g)** adult, **h)** aged, and **i)** D14 FSGS glomerulus. **j)** Line profiles of the thickness distributions of GBM for adult, aged, and D14 FSGS glomeruli. The distributions in **e, f, j, k)** are normalized to have the same area under the curve. All scale bars and distances are shown in pre-expansion units.

The thickness of the BCBM (**Figure 6b-d**) in healthy adult glomeruli was 637 ± 103 nm, with notable variability among individual glomeruli (**Figure 6f**). The BCBM significantly thickened in aged and FSGS glomeruli to 820 ± 128 nm and 851 ± 88 nm, respectively (**Figure 6e**). The thickening in the aged sample primarily occurred in patchy regions at the junction of the Bowman’s capsule and proximal convoluted tubule (PCT, **Figure 6c**), suggesting a potential association between the PCT and the glomerular aging process. In experimental FSGS glomeruli, the thickening occurs specifically at sites of pathological changes, where PECs formed a layered BCBM (**Figure 6d**).

For the GBM (**Figure 6g-i**), measurements were restricted to regions of GBM that are in contact with capillary loops and Bowman’s space only. Additionally, we limited our analysis to regions of the GBM where the surface normal vectors were oriented approximately within the xy-plane (i.e., BM surface normal vectors having an angle of 80-90 degrees with respect to the z-axis). This selection ensured that the GBM, which is ≥200 nm thick, would only be measured along the lateral dimensions where the effective resolution of ∼100 nm would not appreciably broaden the measurement. Due to anisotropy in the microscope’s point spread function, the effective spatial resolution can be as high as ∼350 nm along the axial dimension (**Supplementary Figure S8**). While this orientation criterion excluded parts of the GBM, the large sample size of over 40,000 data points per sample nonetheless covers regions across the entire glomerulus. The thickness of the aged sample significantly increased to 544 ± 46 nm as compared to 297 ± 48 nm in adult samples, while the FSGS samples showed a slight increase to 378 ± 77 nm (**Figure 6j**). No obvious patterns of GBM thickness variation were evident on the GBM for aged or experimental FSGS glomeruli (**Figures 6g-i**).

## Discussion

In this study, we present a novel pipeline for nanoscale three-dimensional imaging, reconstruction, and analysis of whole glomeruli from healthy young adult mice, aged mice, and mice with experimental FSGS.

Our approach offers a novel perspective by examining phenotypic alterations within the whole glomerulus in 3D, thereby providing new insights into glomerular morphology and spatial coordination among structures. Reconstructing glomerular structures in 3D not only validated previously known observations from 2D analyses but also provided more precise measurements and new morphometric details. We classified all major cell types and their total counts within individual glomeruli, observing higher numbers of cells per glomerulus than was reported in prior work possibly as the result of our higher spatial resolution. We also observed that all glomerular cell types undergo hypertrophic enlargement of nuclei during aging and FSGS, not just podocytes as had been previously reported^20^.

Using our high-resolution, whole-glomerulus reconstructions, we were able to perform a novel global thickness analysis of both the BCBM and GBM. Our basement membrane thickness results were slightly higher than previous reports, but this likely results from differences in the measurements. While traditional EM methods use negative stain against tissue sections to determine the distance between adjacent cellular layers as the basement membrane thickness^35,36^, our FLARE stain^18,19^ uses an adapted fluorescent analog of the PAS stain that labels oxidized carbohydrates and may therefore stain a more diffuse or somewhat larger layer of carbohydrates^37,38^. Our analysis was also applied to the entire glomerulus rather than to a small set of thin sections. Despite these differences, our methodology is sensitive to meaningful basement membrane changes. We observed distinctive patterns of thickening of the BCBM in glomeruli from aged (27-29 m.o.) and model FSGS glomeruli at the glomerulus-proximal tubule junction. In contrast, we did not observe clear patterns of thickening for the GBM which instead thickens more globally.

In a recent publication by Miyaki et al.^39^, an electron microscopy (EM) array tomography method was developed to reconstruct whole glomeruli. The approach obtained remarkable data with a wealth of ultrastructural details, including analysis of capillary networks and podocyte foot processes, and in many ways is a complementary assay to our optical approach. While EM array tomography provides higher spatial resolution, it utilizes an involved and lengthy sample processing pipeline that requires specialized instruments and expertise. Our approach on the other hand achieves more modest spatial resolution (∼100 nm) with sample processing procedures that are simpler, faster, and require only commonly available optical microscopes. We further demonstrated the ability to integrate molecular-specific labeling (e.g., cell-specific immunostains) together with our whole glomerulus analysis, which should open numerous future studies including the possibility of studying relationships of key protein or biomarker distributions across whole glomeruli.

Some limitations of our work merit discussion. Our higher cell counts than previous reports could arise from differences in mouse strain or age, or could be influenced by the fact that our work was limited to cortical glomeruli rather than a mix of cortical and juxtamedullary glomeruli. In addition, we did not account for the potential presence of immune cells, although this is unlikely to fully account for the differences in cell numbers that we observed, especially in healthy adult tissues where immune cell presence is typically minimal. Producing these unique whole-glomerulus models is labor-intensive, currently requiring around 3 days of human effort per glomerulus even with the assistance of a machine-learning algorithm. Future improvements, such as a refined algorithm produced with more extensive training data, could potentially expedite this process. In addition, studies focusing on specific compartments or cell types that don’t require segmentation of all structures, would require significantly less analysis, allowing for faster insights tailored to particular research needs.

In summary, we have presented a novel pipeline to image, reconstruct, and quantify the glomerulus as a whole functional unit. By sharing our models and measurements as open digital resources, we aim to support researchers, nephrologists, pathologists, and medical engineers in exploring the complex global relationships among glomerular structures. These models serve as a valuable resource for facilitating future advances. For instance, our models would be suitable as the basis for running realistic fluid mechanics simulations or as templates for bioengineers seeking to fabricate biology-inspired artificial glomeruli by bioprinting. We strongly believe that the biomedical research community benefits greatly from the dissemination of data, results, and tools and we hope that these resources may help the field move forward.

## Acknowledgements

The authors acknowledge funding support from the University of Washington (J.C.V., S.W.); National Heart, Lung and Blood Institute (F31HL168825 to M.W.); National Institute of Diabetes and Digestive and Kidney Diseases (R21DK136026 and U54DK137328 to J.C.V.; R01DK056799, R01DK097598, R01DK126006, R01DK135716, UC2DK126006 to S.J.S.); the Department of Defense (PR180585 to S.J.S.); the NIDDK Innovative Science Accelerator Program (ISAC, www.isac-kuh.org, DK128851 to J.C.V. and S.W.); Washington Research Foundation (Postdoctoral Fellowship to C.P.). We thank the Biology Imaging Core at the University of Washington for their assistance with imaging, and the Keck Imaging Center at the University of Washington for their help with Imaris.

