## Supplementary for "Nanoscale Optical Imaging, Reconstruction, and Spatial Analysis of Whole Mouse Glomeruli"

**1 Department of Chemistry, University of Washington, Seattle, Washington, USA**

**2 Paul G. Allen School of Computer Science & Engineering, University of Washington, Seattle, Washington, USA**

**3 Division of Nephrology, University of Washington,, Seattle, Washington, USA**

**4 Department of Physiology and Biophysics, University of Washington, Seattle, Washington, USA**


**This PDF file includes:**

Supplementary Figure S1-8
Supplementary Tables S1-3
Supplementary Movie S1-4
Supplementary Methods
Supplementary References

**Supplementary Figures**

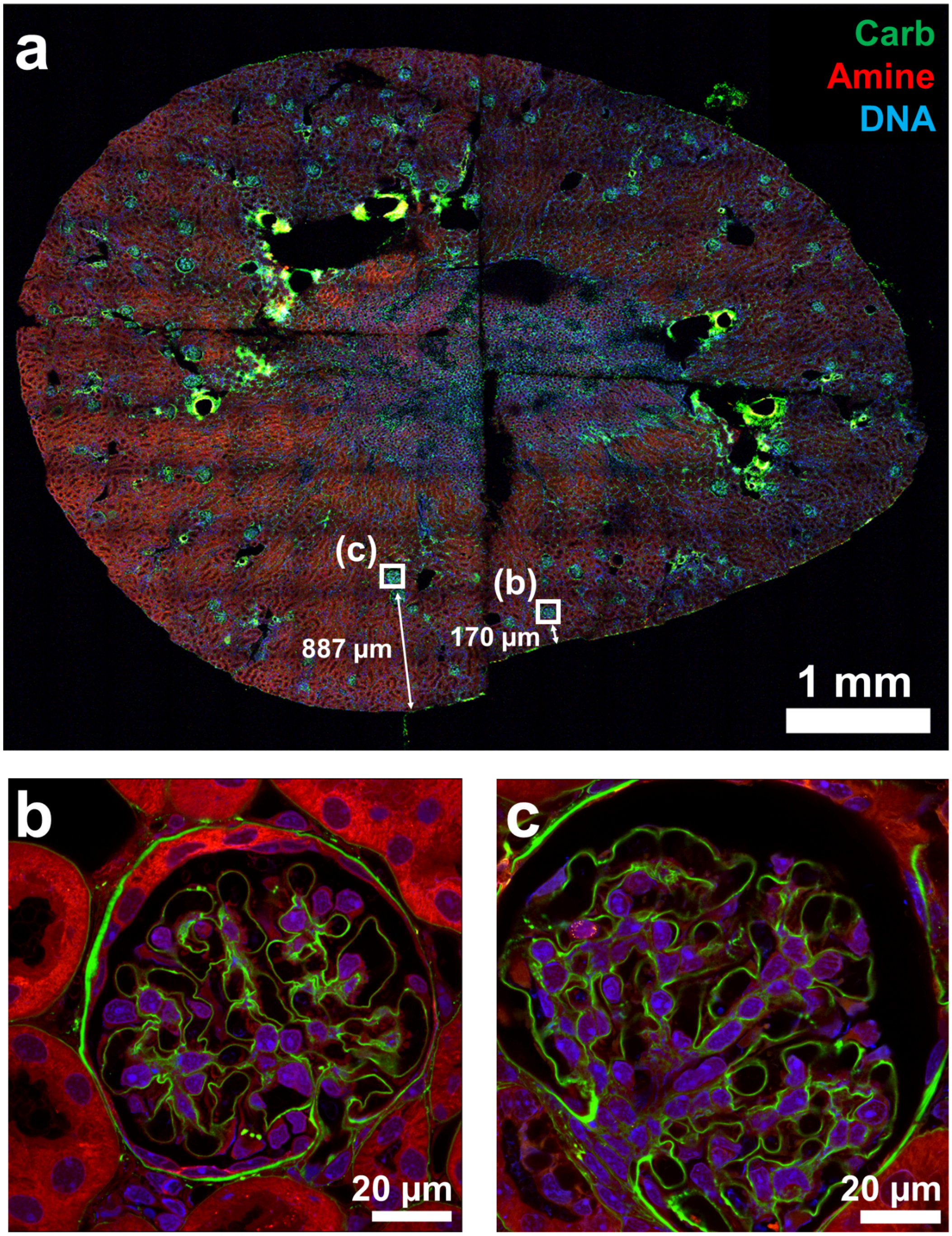


***Supplementary Figure S1***. *Identification and validation of glomeruli location. To validate that the glomeruli acquired in this work were cortical glomeruli, we first imaged* ***a)*** *a map of an entire FLARE-stained expanded kidney section. Zoomed-in views of* ***b)*** *a cortical glomerulus from the superficial region of the kidney section (42µm from the edge) and* ***c)*** *a juxtamedullary glomerulus (219µm from the edge) were imaged using the same imaging center used in the pipeline. Using the same imaging setting, none of the juxtamedullary glomeruli were able to fit in the field of view, confirming that all the glomeruli were cortical. Scale bars are shown in pre-expansion units.*

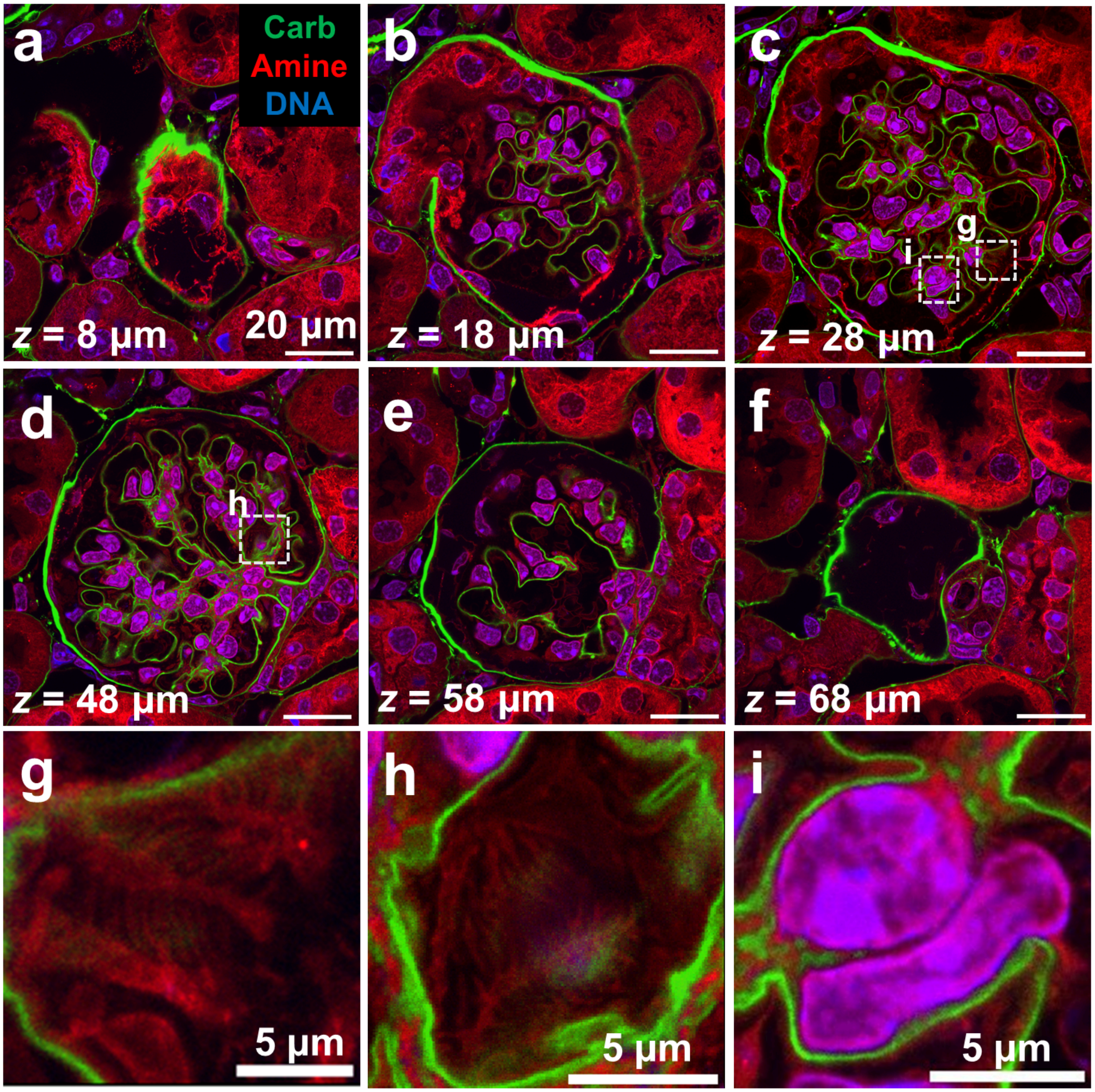


***Supplementary Figure S2***. *A FLARE-stained, hydrogel-expanded mouse glomerulus.* ***a-f)*** *The displayed confocal microscopy optical sections and zoom-in panels* ***g-i)*** *illustrate the level of detail attainable in the imaging approach including* ***g-h)****interdigitated podocyte foot processes, and* ***i)*** *a dense cluster of nuclei within the mesangium. The z-position indicates the position of the optical section within the overall stack. Green=carbohydrates; red=amine/protein, blue=nuclei. Scale bars and z-positions are in pre-expansion units.*

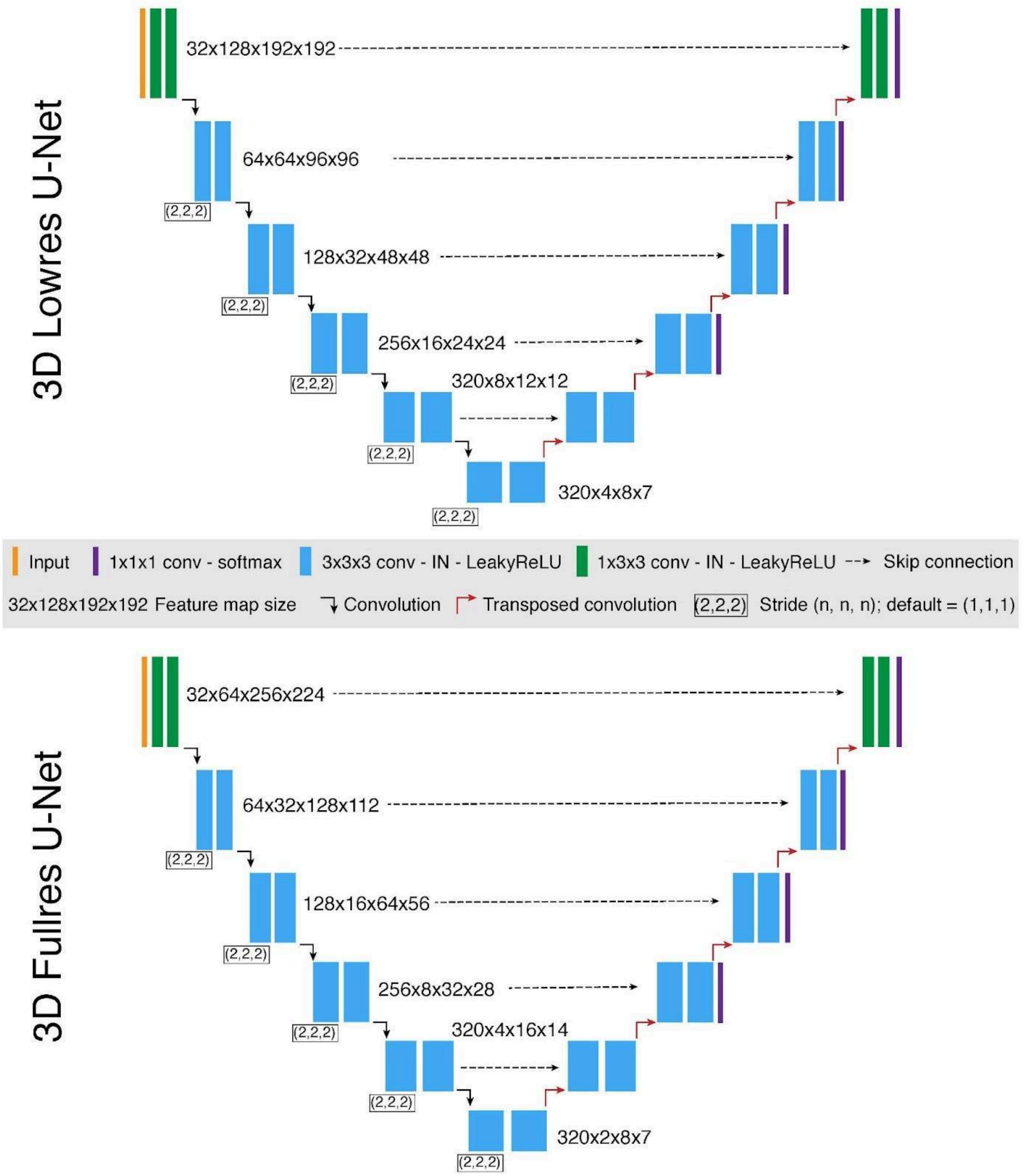


***Supplementary Figure S3***: *Illustration of deep learning-based semi-automated neural network structure. The deep learning framework is composed of a low-resolution 3D CNN-based U-Net and a full-resolution 3D CNN-based U-Net. The low-resolution network takes downsampled glomerular volumes as input to efficiently capture global contextual information, and output coarse predicted segmentation masks. The full-resolution network then takes both the full-sized glomerular volumes and the coarse segmentation masks as input and outputs the refined predicted segmentation masks.*

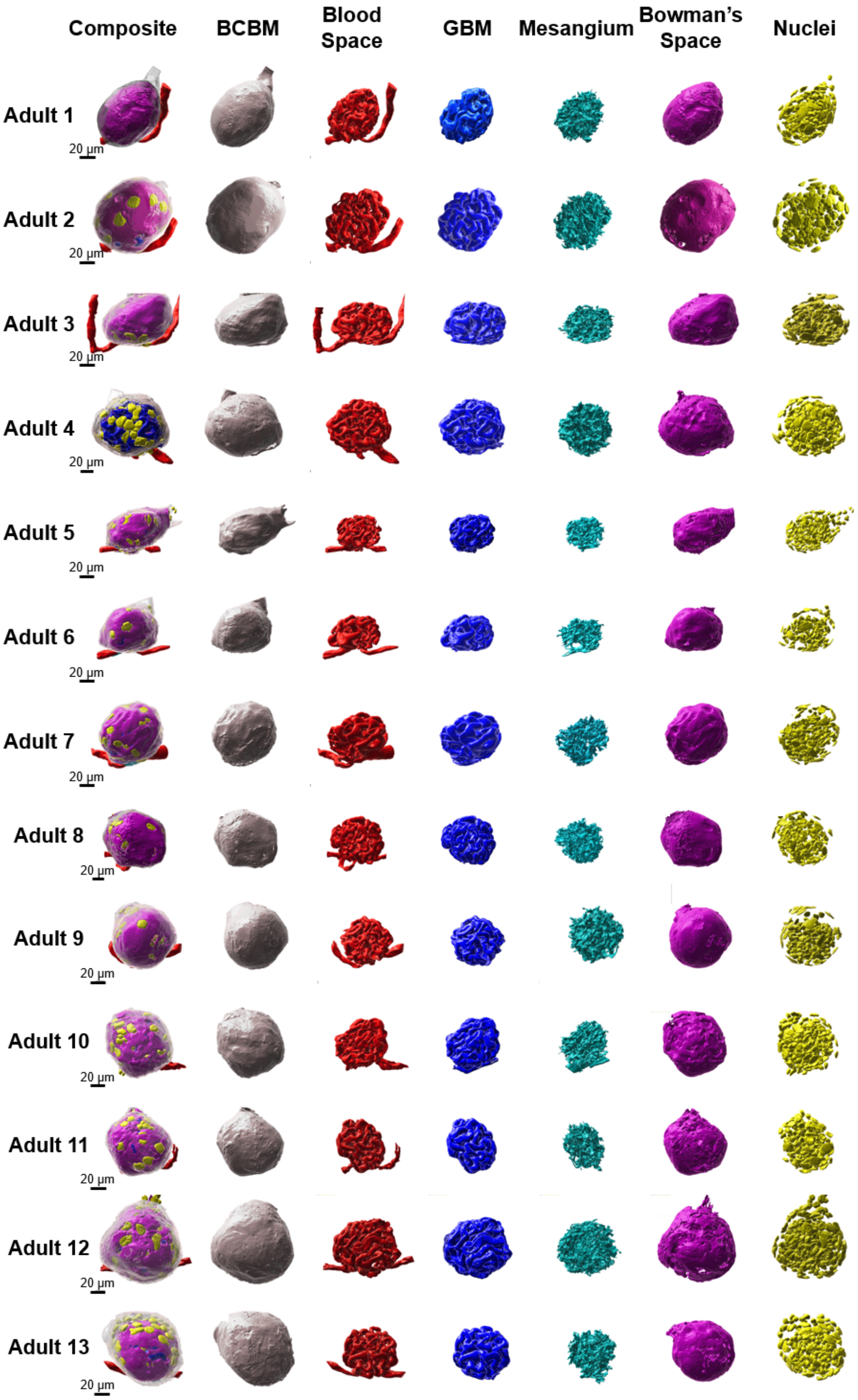
Composite
BCBM
Blood Space
GBM
Mesangium
Bowman's Space
Nuclei
Adult 1
Adult 2
Adult 3
Adult 4
Adult 5
Adult 6
Adult 7
Adult 8
Adult 9
Adult 10
Adult 11
Adult 12
Adult 13
20 µm

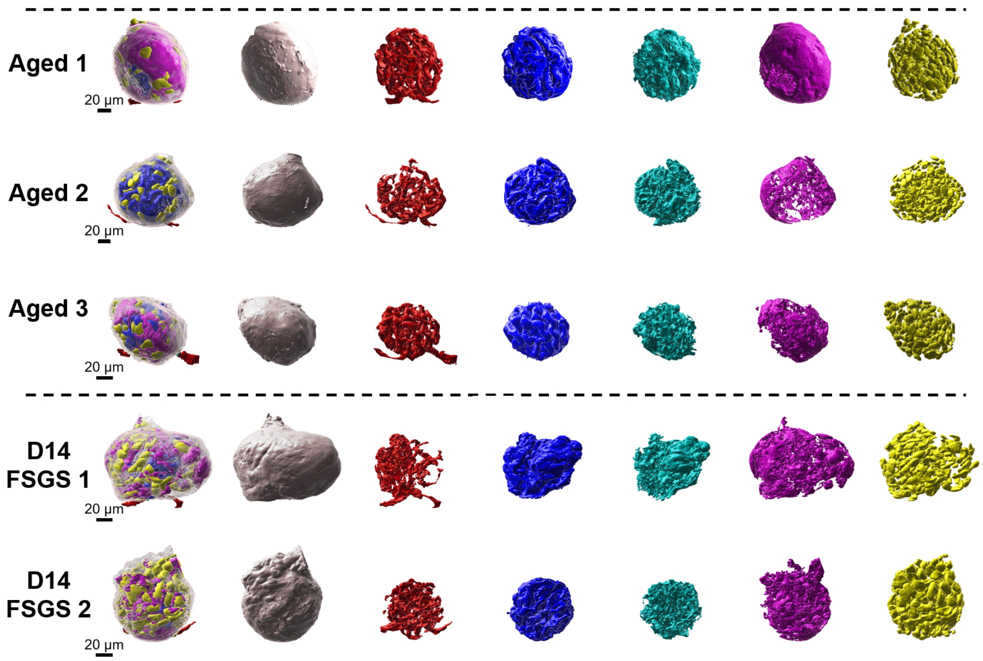


***Supplementary Figure S4***. *Collection of all reconstructed healthy adult, aged, and FSGS mouse glomeruli from this work. All the glomeruli are oriented such that the capillary pole is at the bottom and the urinary pole is at the top. BCBM = grey, Blood space = red, GBM = blue, Mesangium = cyan, Bowman's Space = magenta, Nuclei = yellow. Scale bars are shown in pre-expansion units.*

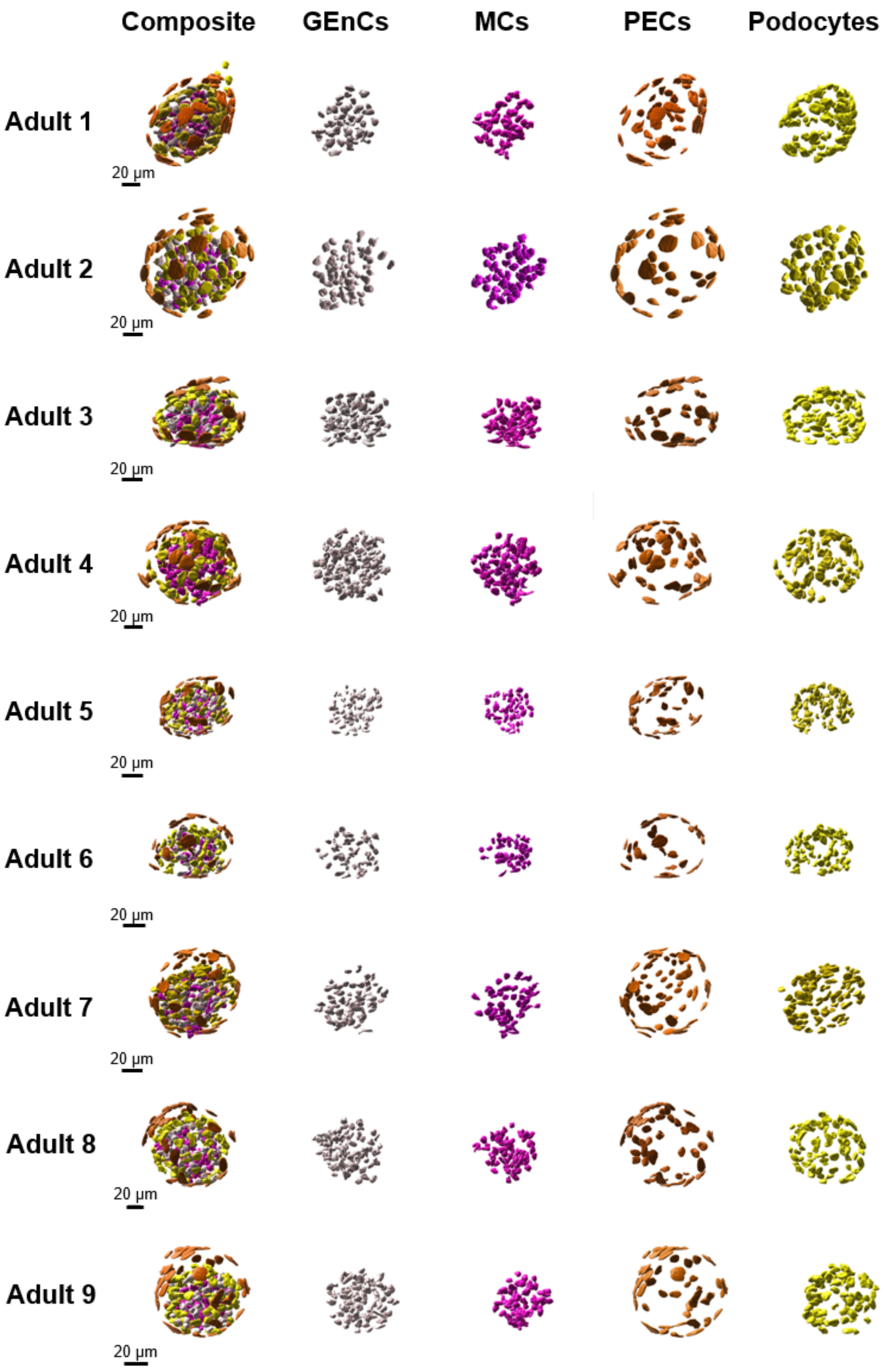
Composite
GEnCs
MCs
PECs
Podocytes
Adult 1
Adult 2
Adult 3
Adult 4
Adult 5
Adult 6
Adult 7
Adult 8
Adult 9
20 µm

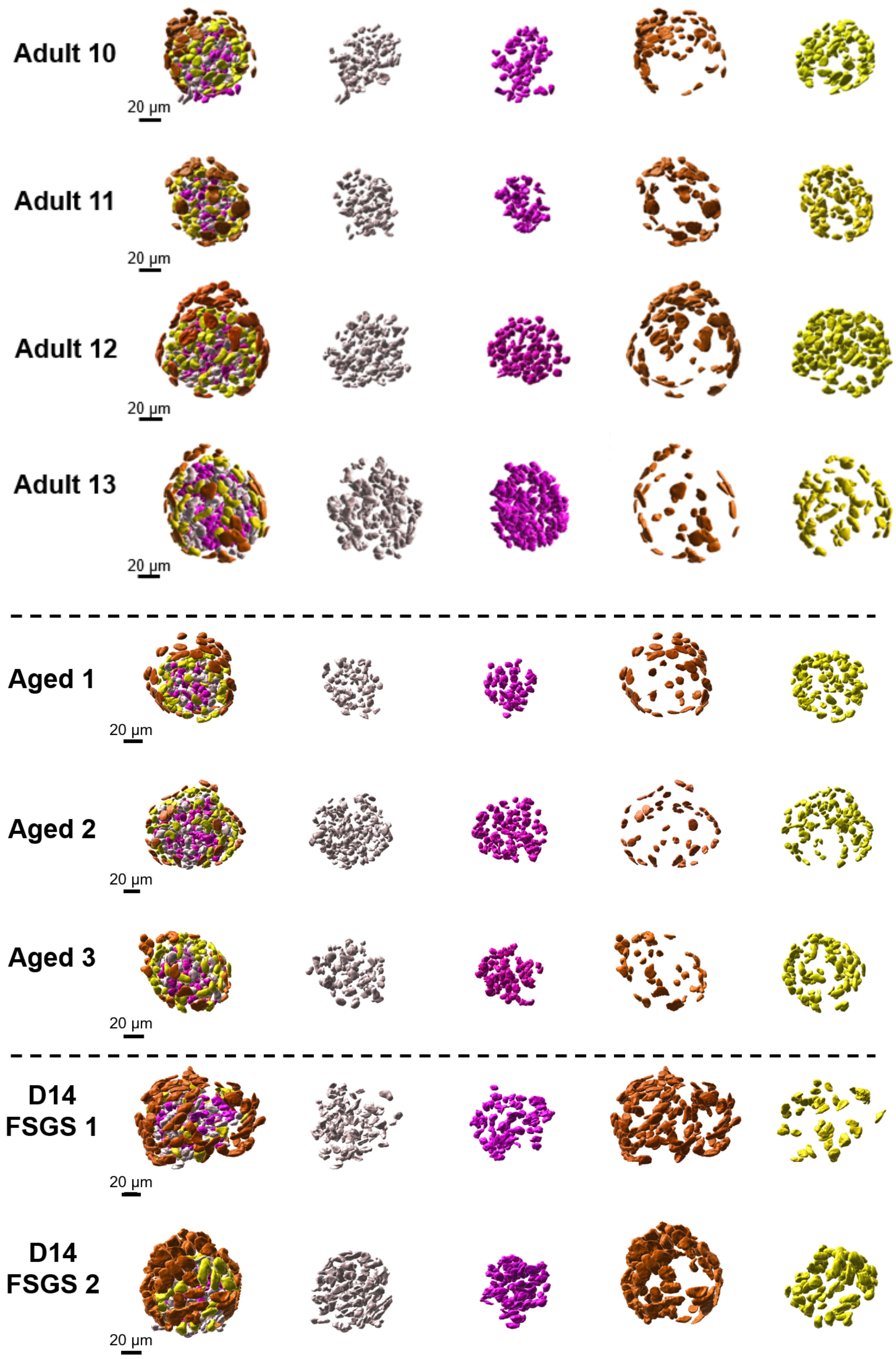


***Supplementary Figure S5***. *Collection of all reconstructed and classified cell nuclei for healthy adult, aged, and FSGS mouse glomeruli. All the images are positioned in the exact same orientation as the glomerular structures in Supplementary Figure 4. GEnCs = grey, MCs = magenta, PECs = blue, Podocytes = yellow. Scale bars are shown in pre-expansion units.*

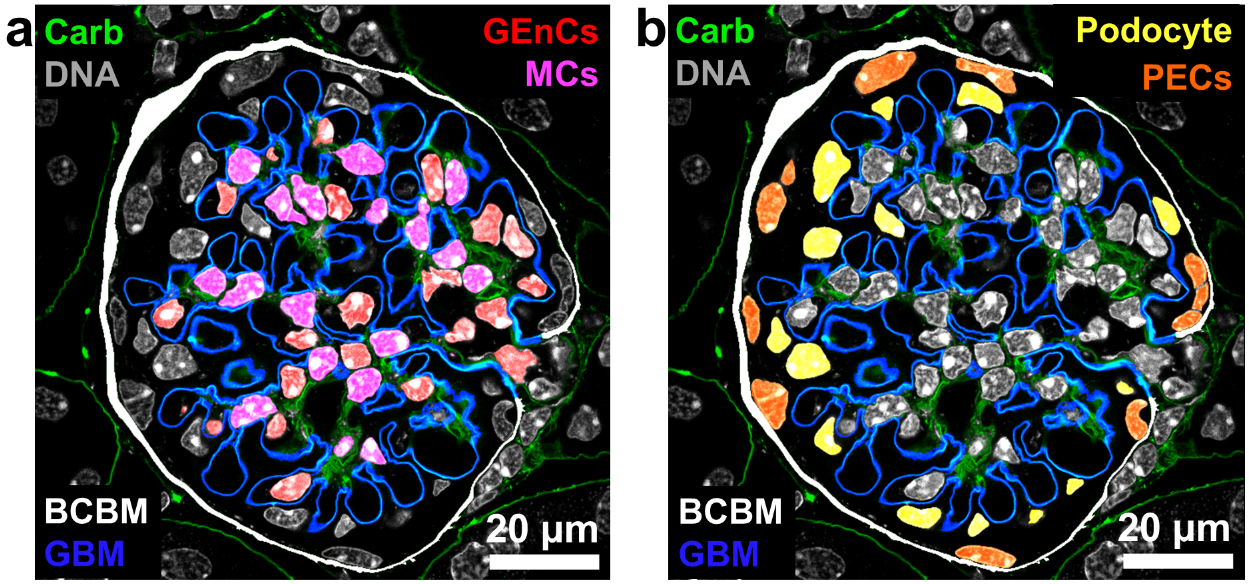


***Supplementary Figure S6***. *Methodology for cell type classification. Segmented labels of GBM (blue), and BCBM (white) are overlaid with the raw carbohydrate channel (green) and raw nuclear channel (grey) to provide guidance on the positioning of the cell nuclei relative to segmented glomerular structures.* ***a)*** *GEnCs and MCs are positioned on the inner side of the GBM (blue), but GEnC nuclei (red) have substantial proximity to blood space (black) while MC nuclei are nearly surrounded by the carbohydrate signal (green) or GBM (blue).* ***b)*** *PEC cell nuclei (orange) are classified as those lining the inner surface of the BCBM (grey), whereas the podocyte nuclei (yellow) reside just outside the GBM (blue). Nuclei were designated as belonging to GEnC, MC, PEC, or podocytes after examining the 3D data set near each nucleus, although the identity of some nuclei is less clear in single-plane views. Scale bars are shown in pre-expansion units.*

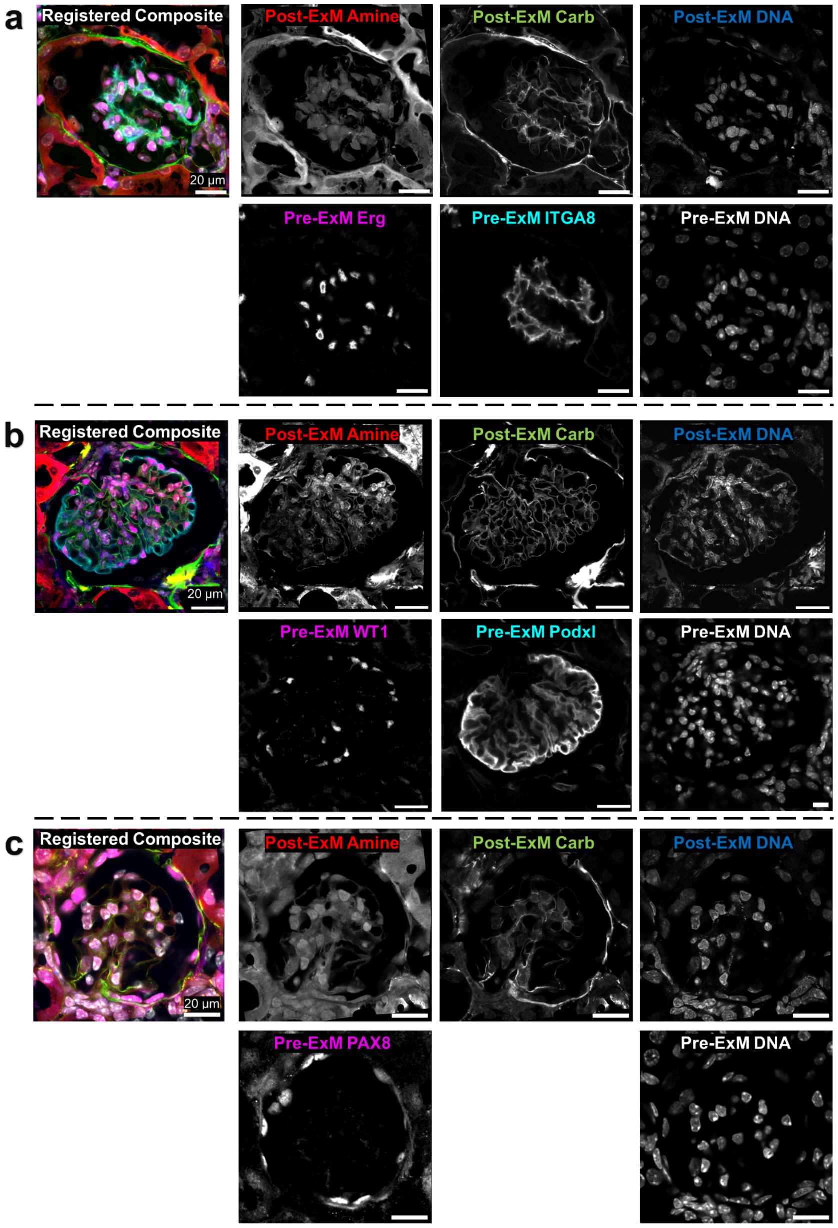


***Supplementary Figure S7****. Validation of cell type classification using pre-ExM immunostains and post-ExM FLARE stains. Mouse kidney tissue sections were initially stained with markers for specific cell types:* ***a)*** *glomerular endothelial cells (Erg) and mesangial cells (ITGA8),* ***b)*** *podocytes (WT1, Podxl), and* ***c)*** *parietal epithelial cells (PAX8). Glomeruli in the unexpanded sections were imaged in a first round. The sections were then processed with expansion microscopy and FLARE staining (that chemically bleaches the prior immunostain signals) and then the same glomeruli were imaged in a second round. Finally, the two rounds of images were registered to produce a single composite data set. Based on this validation methodology, the precision and recall for ExM/FLARE-based cell type classification were determined to be 91-100% (see also* ***Supplementary Table 3****). All scale bars are shown in pre-expansion units.*

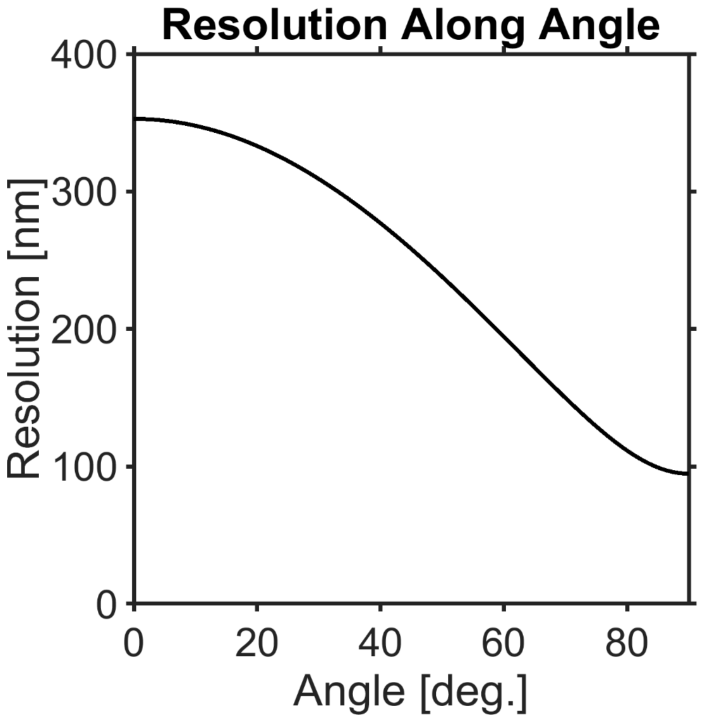


***Supplementary Figure S8***. *Simulation of spatial resolution as a function of angle relative to the ± z dimension. The angle ranges from 0 to 90 degrees, where 90 degrees represents alignment parallel to the xy-plane where the expansion-corrected lateral spatial resolution is ~100 nm, and 0 degrees represents alignment along the ± z-axis where the expansion-corrected axial spatial resolution is ~350 nm. The plot illustrates how the spatial resolution varies between the lateral and axial dimensions due to anisotropy in the point spread function (PSF).*

**Supplementary Table S1**. Average measurements of individual glomerular structures and cell types. All volumes or surface areas are presented in pre-expansion units. Values are presented as mean ± standard deviation.

| | Adult (3-15 m.o.) | Aged (27-29 m.o.) | D14 FSGS (3.5 m.o.) |
|---|---|---|---|
| $V_{Blood}$(×10$^4$ μm$^3$) | 6.76 (±2.45) | 7.73 (±2.91) | 2.66 (±0.08) |
| $V_{Mesangium}$ (×10$^4$ μm$^3$) | 3.60 (±1.48) | 21.9 (±11.7) | 6.39 (±1.13) |
| $V_{GBM}$ (×10$^4$ μm$^3$) | 7.70 (±2.54) | 14.8 (±6.84) | 8.59(±0.96) |
| $V_{BCBM}$ (×10$^4$ μm$^3$) | 4.84 (±1.42) | 10.1 (±3.92) | 9.71 (±1.29) |
| $V_{Nuclei}$ (×10$^4$ μm$^3$) | 3.89 (±1.74) | 9.19 (±3.95) | 8.57 (±0.99) |
| $V_{Bowman's\ Space}$ (×10$^4$ μm$^3$) | 14.3 (±5.12) | 8.97 (±7.28) | 7.45 (±2.92) |
| $V_{Renal\ Corpuscle}$ (×10$^4$ μm$^3$) | 40.2 (±13.7) | 79.8 (±33.3) | 50.9 (±6.46) |
| % $V_{Blood}$ | 17.1 (±3.6) | 10.0 (±1.9) | 5.3 (±0.5) |
| % $V_{Mesangium}$ | 8.8 (±1.4) | 26.1 (±5.2) | 12.5 (±0.6) |
| % $V_{GBM}$ | 19.4 (±2.6) | 18.3 (±1.8) | 16.9 (±0.3) |
| % $_{VBCBM}$ | 12.5 (±2.6) | 13.0 (±1.5) | 19.1 (±0.1) |
| % $V_{Nuclei}$ | 9.4 (±2.3) | 11.5 (±0.7) | 17.1 (±4.1) |
| % $V_{Bowman's\ Space}$ | 36.2 (±6.5) | 10.9 (±6.2) | 14.4(±3.9) |
| $S_{BCBM}$ (×10$^4$ μm$^2$) | 5.27 (±1.1) | 20.5 (±22.3) | 9.55 (±2.50) |
| $S_{GBM}$ (×10$^4$ μm$^2$) | 10.7 (±3.2) | 30.0 (±2.44) | 11.2 (±0.31) |
| Total Cell Count | 319 (±60) | 423 (±87) | 364 (±44) |
| GEnC Count | 92 (±26) | 152 (±53) | 106 (±7) |
| MC Count | 73 (±14) | 133 (±44) | 85 (±19) |
| Podocyte Count | 97 (±22) | 91 (±15) | 54 (±25) |
| PEC Count | 58 (±16) | 46 (±2) | 120 (±7) |
| GEnC% | 28.5 (±3.8) | 35.2 (±6.1) | 29.2 (±1.6) |
| MC% | 22.8 (±1.9) | 31.0 (±5.4) | 23.1 (±2.5) |
| Podocyte% | 30.7 (±5.5) | 22.5 (±7.1) | 14.4 (±5.1) |
| PEC% | 18.1 (±4.3) | 11.3 (±2.4) | 33.3 (±6.0) |
| Nuclear $V_{GEnCs}$ (μm$^3$) | 101.2 (±38.1) | 191.4 (±50.0) | 182.8 (±23.6) |
| Nuclear $V_{MCs}$ (μm$^3$) | 105.7 (±42.3) | 210.3 (±65.1) | 181.0 (±0.2) |
| Nuclear $V_{Podocytes}$ (μm$^3$) | 128.6 (±48.0) | 220.2 (±66.7) | 287.4 (±7.5) |
| Nuclear $V_{PECs}$ (μm$^3$) | 161.3 (±69.3) | 252.7 (±68.0) | 296.9 (±16.5) |

**Supplementary Table S2**. Collective measurements of individual glomerular structures and cell types. All volume and surface area measurements are presented in pre-expansion units.

| | Adult | | | | | | | | | | | | | Aged | | | FSGS | |
|---|---|---|---|---|---|---|---|---|---|---|---|---|---|---|---|---|---|---|
| **Glom** | **1** | **2** | **3** | **4** | **5** | **6** | **7** | **8** | **9** | **10** | **11** | **12** | **13** | **1** | **2** | **3** | **1** | **2** |
| **Sex** | M | M | F | F | M | M | M | F | F | M | M | M | M | F | F | M | M | M |
| **Age** | 6 m | 6 m | 10 m | 10 m | 6 m | 6 m | 6 m | 2 m | 2 m | 4.5 m | 4.5 m | 4.5 m | 4 m | 29 m | 29 m | 27 m | 3.5 m | 3.5 m |
| **Expansion Factor** | 4.0 | 4.0 | 3.7 | 3.6 | 3.9 | 3.9 | 3.9 | 3.3 | 4.0 | 3.8 | 3.9 | 3.9 | 3.9 | 3.5 | 3.5 | 3.7 | 4.0 | 3.9 |
| **$V_{Blood\ Space}$ (×10⁴ μm³)** | 5.2 | 6.2 | 6.2 | 12.3 | 3.2 | 5.2 | 8.5 | 10.9 | 6.3 | 6.2 | 5.3 | 6.3 | 6.0 | 10.5 | 7.9 | 4.7 | 2.7 | 2.6 |
| **$V_{Mesangium}$ (×10⁴ μm³)** | 3.7 | 5.2 | 2.9 | 6.5 | 1.6 | 2.0 | 2.5 | 5.7 | 2.4 | 3.6 | 2.7 | 4.1 | 3.8 | 29.7 | 27.5 | 8.5 | 7.2 | 5.6 |
| **$V_{GBM}$ (×10⁴ μm³)** | 7.4 | 7.6 | 6.5 | 12.8 | 4.7 | 3.8 | 7.6 | 12.1 | 5.7 | 9.0 | 7.0 | 8.2 | 7.5 | 16.6 | 20.5 | 7.2 | 9.3 | 7.9 |
| **$V_{BCBM}$ (×10⁴ μm³)** | 6.3 | 5.4 | 3.9 | 7.8 | 3.7 | 3.3 | 4.3 | 6.8 | 3.5 | 4.6 | 3.9 | 5.4 | 3.8 | 13.4 | 11.3 | 5.8 | 10.6 | 8.8 |
| **$V_{Nuclei}$ (×10⁴ μm³)** | 3.3 | 4.2 | 4.3 | 7.3 | 1.2 | 1.3 | 2.6 | 5.6 | 2.9 | 4.0 | 3.7 | 5.9 | 4.6 | 11.9 | 11.0 | 4.7 | 7.9 | 9.3 |
| **$V_{Bowman's\ Space}$ (×10⁴ μm³)** | 14.3 | 16.6 | 12.6 | 21.2 | 8.2 | 8.7 | 13.8 | 26.5 | 17.0 | 11.9 | 10.0 | 14.0 | 11.4 | 17.3 | 5.9 | 3.7 | 9.5 | 5.4 |
| **$V_{Renal\ Corpuscle}$ (×10⁴ μm³)** | 37.8 | 43.2 | 33.1 | 65.9 | 21.1 | 21.7 | 36.4 | 62.9 | 34.5 | 38.6 | 30.0 | 49.6 | 47.7 | 97.2 | 100.8 | 41.5 | 55.5 | 46.3 |
| **% $V_{Blood}$** | 13.7 | 14.5 | 18.7 | 18.7 | 15.2 | 24.1 | 23.3 | 17.3 | 18.1 | 16.2 | 17.7 | 12.6 | 12.6 | 10.9 | 7.9 | 11.4 | 4.9 | 5.6 |
| **% $V_{Mesangium}$** | 9.9 | 12.1 | 8.9 | 9.8 | 7.4 | 9.4 | 7.0 | 9.0 | 6.9 | 9.2 | 9.0 | 8.3 | 8.0 | 30.6 | 27.3 | 20.5 | 13.0 | 12.1 |
| **% $V_{GBM}$** | 19.7 | 17.6 | 19.5 | 19.5 | 22.5 | 17.7 | 20.9 | 19.2 | 16.4 | 23.3 | 23.4 | 16.6 | 15.8 | 17.1 | 20.3 | 17.4 | 16.7 | 17.1 |
| **% $_{VBCBM}$** | 16.8 | 12.6 | 11.8 | 11.8 | 17.4 | 15.3 | 12.0 | 10.9 | 10.3 | 12.0 | 12.9 | 10.8 | 7.9 | 13.8 | 11.2 | 13.9 | 19.2 | 19.0 |
| **% $V_{Nuclei}$** | 8.6 | 9.6 | 12.9 | 11.0 | 5.8 | 5.8 | 7.0 | 8.9 | 8.3 | 10.3 | 12.5 | 11.8 | 9.6 | 12.3 | 10.9 | 11.2 | 14.2 | 20.0 |
| **% $V_{Bowman's\ Space}$** | 37.9 | 38.3 | 38.1 | 32.1 | 38.7 | 39.9 | 37.9 | 42.2 | 49.3 | 30.9 | 33.4 | 28.3 | 23.8 | 17.8 | 5.8 | 9.0 | 17.2 | 11.6 |
| **$S_{BCBM}$ (×10⁴ μm²)** | 5.0 | 5.6 | 4.7 | 7.4 | 3.7 | 3.8 | 4.8 | 7.3 | 4.7 | 5.3 | 4.6 | 6.0 | 5.6 | 46.1 | 9.7 | 5.7 | 11.3 | 7.8 |
| **$S_{GBM}$ (×10⁴ μm²)** | 10.8 | 10.6 | 11.2 | 18.2 | 6.4 | 6.6 | 9.8 | 13.5 | 8.4 | 9.3 | 9.2 | 14.1 | 11.8 | 21.9 | 22.1 | 10.5 | 10.9 | 11.4 |
| **Total Cell Count** | 282 | 264 | 294 | 378 | 291 | 234 | 301 | 312 | 310 | 383 | 317 | 466 | 317 | 479 | 467 | 322 | 333 | 395 |
| **GEnC Count** | 67 | 66 | 86 | 140 | 76 | 57 | 77 | 98 | 102 | 108 | 93 | 139 | 87 | 180 | 186 | 91 | 101 | 111 |
| **MC Count** | 63 | 70 | 74 | 94 | 66 | 55 | 62 | 73 | 63 | 88 | 66 | 101 | 68 | 178 | 130 | 90 | 71 | 98 |
| **Podocyte Count** | 105 | 86 | 92 | 89 | 102 | 86 | 88 | 87 | 54 | 112 | 100 | 152 | 108 | 75 | 103 | 96 | 36 | 71 |
| **PEC Count** | 47 | 42 | 42 | 55 | 47 | 36 | 74 | 54 | 91 | 75 | 58 | 74 | 54 | 46 | 48 | 45 | 125 | 115 |
| **GEnC %** | 24 | 25 | 29 | 37 | 26 | 24 | 26 | 31 | 33 | 28 | 29 | 30 | 27 | 38 | 40 | 28 | 30 | 28 |
| **MC%** | 22 | 27 | 25 | 25 | 23 | 24 | 21 | 23 | 20 | 23 | 21 | 22 | 21 | 37 | 28 | 28 | 21 | 25 |
| **Podocyte%** | 37 | 33 | 31 | 24 | 35 | 37 | 29 | 28 | 17 | 29 | 32 | 33 | 34 | 16 | 22 | 30 | 11 | 18 |
| **PEC%** | 17 | 16 | 14 | 15 | 16 | 15 | 25 | 17 | 29 | 20 | 18 | 16 | 17 | 10 | 10 | 14 | 38 | 29 |
| **GEnC Nuclear V (μm³)** | 93.3 | 138.9 | 125.9 | 160.5 | 32.4 | 48.0 | 74.4 | 152.2 | 81.8 | 86.8 | 98.0 | 106.6 | 116.9 | 221.4 | 219.2 | 133.7 | 166.0 | 199.5 |
| **MC Nuclear V (μm³)** | 98.8 | 133.5 | 131.3 | 179.4 | 31.2 | 44.9 | 69.4 | 160.7 | 85.8 | 91.6 | 110.2 | 114.0 | 123.7 | 248.2 | 247.5 | 135.1 | 180.8 | 181.2 |
| **Podocyte Nuclear V (μm³)** | 118.8 | 157.6 | 154.1 | 209.4 | 47.7 | 56.2 | 90.0 | 184.2 | 168.9 | 96.1 | 119.0 | 123.6 | 146.6 | 281.4 | 230.1 | 149.1 | 292.7 | 282.1 |
| **PEC Nuclear V (μm³)** | 162.8 | 227.2 | 189.7 | 269.1 | 61.7 | 71.2 | 103.1 | 243.0 | 64.2 | 152.9 | 156.0 | 181.9 | 213.7 | 308.5 | 272.7 | 177.0 | 308.5 | 285.2 |

**Supplementary Table S3**. Validation result for classification of glomerular cell type. Precision = (True positive)/(True positive + false positive). Recall = (True positive)/(True positive + False negative).

| | **Glomerular Endothelial Cell** | **Mesangial Cell** | **Podocyte** | **Parietal Epithelial Cell** |
|---|---|---|---|---|
| **True Positive** | **192** | **139** | **127** | **130** |
| **False Positive** | **17** | **11** | **2** | **1** |
| **False Negative** | **18** | **0** | **11** | **2** |
| **Precision** | **91.9%** | **92.7%** | **98.4%** | **99.2%** |
| **Recall** | **91.4%** | **100.0%** | **92.0%** | **98.5%** |

**Supplementary Movies**

**Supplementary Movie 1**. *Image stack data (first half of movie) and segmented rendering (second half of movie) for the same adult glomerulus as shown in **Figure 2a**. Some white colors in segmented nuclei are due to the overlap between yellow (nuclei) and cyan (mesangium). Scale bars: 20 µm in pre-expansion units.*

**Supplementary Movie 2.** *Image stack data (first half of movie) and segmented rendering (second half of movie) for the same aged glomerulus as shown in **Figure 2b**. Some white colors in segmented nuclei are due to the overlap between yellow (nuclei) and cyan (mesangium). Scale bars: 20 µm in pre-expansion units.*

**Supplementary Movie 3**. *Image stack data (first half of movie) and segmented rendering (second half of movie) for the same D14 FSGS glomerulus as shown in **Figure 2c**. Some white colors in segmented nuclei are due to the overlap between yellow (nuclei) and cyan (mesangium). Scale bars: 20 µm in pre-expansion units.*

**Supplementary Movie 4**. *3D view of the same reconstructed nuclei of individual cell types and corresponding glomerular compartments as shown in **Figure 5a**. Note that the efferent arteriole exceeds the field of view and was partly truncated. Scale bars: 20 µm in pre-expansion units.*

### Supplementary Methods

#### Reagents

Reagents for hydrogel polymerization, based on the Magnified Analyais of the Proteome (MAP)[1] hydrogel protocol for polymerization, were purchased as follows: 40% (w/v) acrylamide (AA; Bio-Rad Laboratories, cat. no. 161-0140), 2% (w/v) bis-acrylamide (BA; Bio-Rad Laboratories, cat. no. 161-0142), Sodium acrylate (SA; Sigma-Aldrich, cat. no. 408220), VA-044 (Thermo Fisher Scientific, cat. no. NC0471397), bovine serum albumin (BSA; Rockland Immunochemicals Inc, item no. BAS-50). Reagents for denaturation were purchased as follows: Sodium dodecyl sulfate (SDS; Sigma-Aldrich, cat. no. L3771), Tris base (Tris; Fisher Scientific, cat. no. BP152-500), Sodium chloride (NaCl; Fisher Scientific, cat. no. 271). Reagents for FLARE staining were purchased as follows: 10× phosphate-buffered saline, pH 7.4 (PBS; Fisher Bioreagents, cat. no. L-5400), Sodium azide (NaN3; Fisher Scientific, cat. no. S227I), Sodium acetate, anhydrous (NaOAc; Fisher Scientific, cat. no. S209), 4-Morpholineethanesulfonic acid (MES; Sigma-Aldrich, cat. no. M8250), NaIO4 (Sigma-Aldrich, cat. no. 311448), Sodium cyanoborohydride ($NaCNBH_3$; Sigma-Aldrich, cat. no. 156159), ATTO-TEC GmbH, cat. no. AD565, ATTO 647N NHS ester (AT647N-NHS; Sigma-Aldrich, cat. no. 18373), ATTO 565 hydrazide ($AT565\text{-}NHNH_2$, Hoechst 33258 (Sigma-Aldrich, cat. no. B2883). Triton X100 (61-2754) and Poly-L-lysine (P8920) were purchased from Sigma-Aldrich. Primary antibodies were purchased as follows: Rabbit anti-Erg (Abcam Inc., cat. no. ab214341), Goat anti-integrin alpha-8 (Fisher Scientific, cat. no. PA5-47572), Rabbit anti-Wilms Tumor Protein antibody (Abcam Inc., cat. no. ab89901), Goat anti-podocalyxin (R&D Systems, cat. no. AF1556), Rabbit anti-Pax8 (Proteintech, cat. no. 10336-1-AP). Secondary antibodies and fluorophores were purchased as follows: Donkey anti-rabbit IgG (Jackson ImmunoResearch Lab Inc, cat. no. 711-005-152), Donkey anti-goat IgG (Jackson Immuno Research Lab Inc, cat. no. 705-005-147), ATTO 488 NHS ester (Millipore Sigma, cat. no. 41698-1MG-F), Alexa Fluor 568 NHS ester (Lumiprobe, cat. no. 44820).

#### Preparation of fluorophore-conjugated secondary antibodies

Fluorophores were conjugated to secondary antibodies in-lab by first mixing ~40 μl of unconjugated secondary antibodies at (~1.3 mg/ml), 10 μl of 1M sodium bicarbonate (~pH 9.3), and 3 μg of NHS dye from a dye stock in anhydrous DMSO (~2 μg/μl). The bioconjugation reaction was allowed to proceed for 5-10 min at room temperature. After the reaction, a NAP-5 size exclusion column (Fisher Scientific, cat. no. 45-000-152) was used for purification. The column was equilibrated with ~10 ml PBS. The reaction mixture was then loaded onto the column followed by ~600 μl of PBS. Another 300 μl of PBS was then added to the column and the eluent was collected. The eluent absorbance was measured by UV/Vis absorption spectroscopy in order to determine the antibody concentration and dye-to-protein ratios (typically 200 μg/ml and 5-8 dye per antibody).

**Immunostaining of kidney tissue samples**

100 µm thick mouse tissue sections were immunostained using the following protocol. Tissue sections were first incubated in blocking/permeabilization buffer (3% BSA and 0.1% Triton X100 in PBS) overnight at 4°C. The sections were then incubated with primary antibodies at 3 µg/ml diluted in blocking/permeabilization buffer overnight at 4°C. The sections were then washed with blocking/permeabilization buffer three times at room temperature, 10 min each. The sections were then incubated with fluorophore-conjugated secondary antibodies at 5 µg/ml diluted in blocking/permeabilization buffer overnight at 4°C. The sections were then washed with blocking/permeabilization buffer three times at room temperature, 10 min each.

**Kidney tissue gelation, denaturation, and expansion**

100 µm thick kidney sections were first incubated in the MAP monomer solution [20% (w/w) AA, 10% (w/v) SA, 0.05% (w/w) BA, and 4% (v/v) PFA] for 24 hours at 4ºC. The section was then placed on a rectangular #1.5 coverslip and gently spread flat using a paintbrush. Two stacked smaller pieces of #1.5 coverslips were placed on either end of the sample to act as spacers. A gelation solution was prepared by mixing 99 µl of pre-warmed MAP monomer solution with 1 µl of 10% (w/v) VA-44. A drop of the gelation solution (10-15 µl) was pipetted onto the tissue surface and covered with another piece of rectangular #1.5 coverslip to create a gelation chamber. The gelation chamber is then placed into a glass chamber and purged with sufficient nitrogen gas before closing the lid. Gelation was conducted for 2-2.5 hours at 45ºC. After gelation, the gel is peeled off and placed in a scintillation vial containing denaturation solution [200 mM SDS, 200 mM NaCl, 50 mM Tris]. The denaturation was performed under 70ºC for 18-24 hours and 95ºC for another 18-24 hours. The gel was washed with PBST solution three times, 30 minutes each, and one more wash with 1× PBS/Azide for another 30 minutes.

**FLARE staining protocol**

Gelled tissue samples were stained in the order of carbohydrates, amines, and DNA. Gelled and denatured samples were first oxidized with 2 ml of 20 mM $NaIO_4$ in NaOAc/NaCl buffer for 1 hour at 37ºC, protected from light with constant shaking, followed by three washes with NaOAc buffer for 10 minutes each. After oxidation, samples were then reacted in 1mL of 5 µg/mL AT565-Hydrazide in NaOAc for 3 hours at room temperature. The samples were finally reduced by the addition of 20 µl of 5 M NaCNBH4 in NaOAc buffer for 30 minutes at room temperature. For staining amines, the samples were incubated with 2 ml of 5 µg/ml AT647N-NHS in MES buffer at pH 6 for 6 hours at room temperature. For DNA labeling, samples were stained with 1mL of 5 µg/ml Hoechst 33258 in 1× PBS for 45 minutes at room temperature, followed by sufficient washes by 1× PBS. All the above procedures were performed under the protection of light with constant shaking. Lastly, the samples were placed in a large quantity of deionized water for complete expansion overnight.

### Fluorescence image acquisition

The MAP-FLARE sample was fixed onto a poly-lysine-coated cover glass to prevent drifting during imaging. Next, the sample was imaged using a scanning confocal microscope (Nikon A1R HD25 laser scanning confocal microscope) at the University of Washington Biology Imaging Center. A 40× water immersion objective lens (Apochromat Lambda S LWD 40× water, NA 1.15, WD 0.59–0.61 mm) was used and ~0.7 µm z-steps were taken for the 3-dimensional stack with concurrent measurements of the three FLARE labels. There are roughly 12,000 glomeruli within a mouse kidney[2,3], however, only 3-5 whole, intact glomeruli were accessible per 100 µm kidney section due to both the working distance of the objective and positioning of the glomeruli within the tissue sections. As a result, a compromise on the resolution had to be taken considering the imaging time and the working distance of the objective. The ~300 nm physical lateral resolution of the objective lens coupled with ~4× physical expansion could in principle achieve ~75 nm effective spatial resolution, but due to limitations of the number of pixels accessible to the fast galvo system on the confocal microscope, we opted to use a ~200 nm lateral physical pixel size that could achieve a sufficiently large field of view to capture whole glomeruli. The lateral pixel size, when accounting for ~4× physical expansion, corresponds to ~50 nm pixels in pre-expansion units and achieves an overall Nyquist-limited lateral resolution of ~100 nm. The axial step size of ~700 nm also slightly undersamples the ~1.2 µm physical axial resolution of the objective lens, and together with the ~4× physical expansion achieved an overall axial spatial resolution of ~350 nm. Together these settings enabled 3-channel detection within a ~1 hour period.

### Manual glomerular segmentation

Structure segmentation and identification were all done using ImageJ software using the following methods:

1) Full three-channel FLARE stained raw data stacks were first split into carbohydrate, amine, and DNA as separate stacks. The noise from the carbohydrate stack was first cleaned up using *Process->Filters->Gaussian Blur 3D* with a sigma of 3. For noisier data sets, a 3D median filter was instead applied using *Process->Filters->Median 3D* with a radius of 3-5 in all directions. These filtering steps eliminated "salt & pepper" noise but retained most of the signal on the capsule. The resulting stack was then binarized using *Process->Binary->Make Binary*. For the binary method, using either "Li", or "mean" gave the best result in keeping most of the capsule signal while minimizing other signals.

2) Careful manual corrections were done along the BCBM to connect disconnected bits along the BCBM using the *Paintbrush Tool*. After correction, the *Paintbrush* tool was then used to erase all the other signals aside from the capsule.

3) The GBM was segmented using the same strategy as the BCBM.

4) Nuclei were segmented by manually tracing all the cell nuclei periphery in the DNA stack using the *Paintbrush* tool and then filled using the *Process->Binary->Fill Holes* tool.

5) The blood space was identified by first filling the segmented GBM using the *Process->Binary->Fill Holes* tool. Then the segmented GBM, nuclei,, and binarized amine channel were subtracted from the filled GBM using the *Process->Image Calculator* tool.

6) The Bowman's space was identified by first filling the segmented BCBM using the *Process->Binary->Fill Holes* tool. Then the segmented GBM, BCBM, nuclei, blood space, and binarized amine channel were then subtracted from the filled BCBM using *Process->Image Calculator* tool.

7) Lastly, the mesangium was segmented by filling the GBM using the *Process->Binary->Fill Holes* tool. Segmented GBM, blood space, and glomerular endothelial cells were then subtracted from the filled GBM result, using the *Process->Image Calculator* tool.

**Semi-automated glomerular segmentation**

Initially, we manually segmented five glomeruli as described above. These segmented structures are used as the foundation for training a neural network model.

Using the relatively small set of manually segmented data, we trained a neural network model to assist with the segmentation of structures in additional glomeruli. Although this did not produce a fully automated algorithm for glomerulus segmentation, it significantly reduced the human time required.

To segment the GBM, and BCBM, we employed the cascaded nn-UNet 3D framework[4]. This framework consists of two main stages: 3DCNN_lowres, and 3DCNN_fullres, which work together to perform both global level and fine-grained segmentation. 3DCNN_lowres processes the glomerulus at a downsampled resolution to capture global contextual information and outputs a coarse segmentation mask for further refinement. It is particularly effective at segmenting the GBM and BCBM due to its ability to provide coarse segmentation masks while reducing computational complexity. To refine the coarse segmentation, the 3DCNN_fullrest network operates at full resolution, using the original image along with the upsampled coarse masks. This step ensures that local details are captured (that may have been lost during first step downsampling) and is used to generate the final segmentation predictions. Both networks are based on an encoder-decoder architecture with residual connections[5] and skip connections[6], improving the flow of information through the network and increasing training efficiency. Each network was trained using a combination of cross-entropy loss and Dice loss[7], which optimizes segmentation accuracy by combining voxel-wise classification and overlap-based metrics.

For training, we used the first 5 manually segmented glomerular volumes and trained both networks for 20 epochs, with a batch size of 2. Z-score normalization and pixel-spacing-based downsampling were used as preprocessing steps. The training time was less than 1 hour for 3DCNN_lowres and around 12 hours for 3DCNN_fullres on a 16GB NVIDIA A6000 GPU. After training, the raw glomerulus volume was downsampled and fed into the 3DCNN_lowres model to get the downsampled approximate mask. Then, the approximate mask was upsampled and combined with the raw volume before being fed into the 3DCNN_fullres network to get the final prediction. The code for training and segmentation model is available at Zenodo repository (https://doi.org/10.5281/zenodo.13989435)

The overall prediction process for each raw glomerulus data set is less than 30 minutes, and the post-prediction manual refinement took 2-3 days per glomerulus, on average, which is much less than the original approximate 4 weeks for complete manual segmentation for a glomerulus.

The trained neural network model produced initial predictions with three channels: GBM, BCBM, and nuclei. To ensure the accuracy of these predictions, we manually reviewed and corrected the results for any discrepancies or errors using the manual methods described.

For the segmentation of the remaining glomerular structures—namely, blood space, Bowman's space, and the mesangium, we employed the same manual segmentation methods above.

**Glomerular structures reconstruction and analysis**

The segmented five-channel stack was imported to Imaris (Bitplane AG, Zurich, Switzerland). Each individual structure was reconstructed using the surface creation tool on Imaris. For specific instructions on the surface creation tool, see Imaris Reference Manual (https://imaris.oxinst.com/).

**Pre- & post-expansion image registration**

Correlative registration of pre-expansion immunostain and post-expansion FLARE samples was performed as described previously using Elastix[8]. Briefly, the nuclear channels of pre- and post-expansion images of the same glomerulus were first roughly aligned using a rigid linear similarity transformation. The same transformation parameters were then applied to the post-expansion carbohydrate and amine channels of the same stack. Next, a non-rigid B-spline transformation was applied to the post-expansion image to match the pre-expansion image. Again, the same transformation parameters were then applied to the other two channels in order to produce a single composite of the pre-expansion immunostains and the post-expansion FLARE signal.

**Cell type validation**

To validate glomerular cell type classification, mouse kidney sections were first immunostained against markers for each individual cell type using the immunostain protocol listed above. An overall map of the stains was imaged using a confocal microscope at ~1.7 µm resolution. Several glomeruli per cell type were imaged at ~0.2 µm resolution, and their positions within the map were noted. The stained and imaged sections were subsequently processed with expansion microscopy and FLARE staining (which bleach the immunostain fluorophores). The same glomeruli were located on the expanded and FLARE stained samples and then imaged by confocal microscopy at high spatial resolution (e.g., 100 nm effective lateral spatial resolution). Pre- & post-ExM images were registered as described above. During cell-type validation data analysis, cell types were first assigned manually by only looking at post-ExM FLARE stained images. These assigned cell types were then validated using the pre-ExM cell-specific immunostains to evaluate the precision and recall of the classification.

Precision and recall are defined as below:

| | Immunostain | FLARE |
|---|---|---|
| True + (TP) | o | o |
| False + (FP) | × | o |
| False – (FN) | o | × |

| | Precision | Recall |
|---|---|---|
| Definition | Of all predicted positives (using FLARE), how many are really positive (using immunostain) | Of all real positive cases (using immunostain), how many are predicted positive (using FLARE) |
| Mathematical Definition | $\frac{TP}{TP + FP}$ | $\frac{TP}{TP + FN}$ |
